# Rab3 mediates cyclic AMP-dependent presynaptic plasticity and olfactory learning

**DOI:** 10.1101/2023.12.21.572589

**Authors:** Divya Sachidanandan, Aishwarya Aravamudhan, Achmed Mrestani, Jana Nerlich, Marius Lamberty, Natalie Hasenauer, Nadine Ehmann, Dennis Pauls, Teresa Seubert, Isabella Maiellaro, Mareike Selcho, Manfred Heckmann, Stefan Hallermann, Robert J. Kittel

## Abstract

Presynaptic forms of plasticity occur throughout the nervous system and play an important role in learning and memory but the underlying molecular mechanisms are insufficiently understood. Here we show that the small GTPase Rab3 is a key mediator of cyclic AMP (cAMP)-induced presynaptic plasticity in *Drosophila*. Pharmacological and optogenetic cAMP production triggered concentration-dependent alterations of synaptic transmission, including potentiation and depression of evoked neurotransmitter release, as well as strongly facilitated spontaneous release. These changes correlated with a nanoscopic rearrangement of the active zone protein Unc13A and required Rab3. To link these results to animal behaviour, we turned to the established role of cAMP signalling in memory formation and demonstrate that Rab3 is necessary for olfactory learning. As Rab3 is dispensable for basal synaptic transmission, these findings highlight a molecular pathway specifically dedicated to tuning neuronal communication and adaptive behaviour.

## MAIN TEXT

Various classic examples of synaptic plasticity involve functional changes at the presynapse that depend on cAMP second messenger signalling^1,2^. It is, however, not well understood which molecular processes are modified by the cAMP pathway during presynaptic plasticity. The GTPase Rab3 is associated with synaptic vesicles^3,4^ and is involved in short- and long-term synaptic plasticity in vertebrates^5–7^ and invertebrates^8,9^. Rab3 has been linked to synaptic vesicle mobilisation, priming, and late stages of the exocytic process^7,10,11^. Moreover, *Drosophila* Rab3 controls the molecular composition of the presynaptic active zone (AZ), the site of neurotransmitter release from synaptic vesicles^8,9^. However, the role of Rab3 in cAMP-dependent presynaptic plasticity remains elusive. This is at least in part due to experimental difficulties of precisely manipulating cAMP dynamics *in situ* and the complex situation created by four Rab3 paralogs expressed in the mammalian brain (Rab3A-D). Here we focus on the single *Drosophila* ortholog and combine optogenetics with modern imaging techniques. Our results show that Rab3 is required for presynaptic plasticity and memory formation by mediating a cAMP-dependent enhancement of synaptic vesicle release.

### cAMP-triggered presynaptic plasticity requires Rab3

To study cAMP-induced presynaptic plasticity we exposed the glutamatergic larval neuromuscular junction (NMJ) to the adenylyl cyclase agonist forskolin (FSK) and recorded excitatory postsynaptic currents (EPSCs) evoked at low stimulation frequency (0.2 Hz; **Fig. 1a**). Bath application of FSK significantly increased EPSC amplitudes over the time course of 20 minutes (**Fig. 1b,c**), whereas the frequency of spontaneously occurring single synaptic vesicle fusion events (miniatures) remained constant and their amplitude decreased slightly (**Fig. 1d,e**). Consistent with previous results at the *Drosophila* NMJ^12^, it follows that FSK increases the quantal content, i.e. the number of synaptic vesicles released per action potential (0 min: 108 ± 9.4, 20 min: 170 ± 9.8 SEM, n=13 NMJs, p<0.0001 paired t-test). In *Drosophila* null mutants of *rab3* (*rab3^rup^*), a decrease in NMJ AZ number is accompanied by an increase in average AZ size and release probability, resulting in normal basal synaptic transmission (**Extended Data Table**)^8,9^. Upon FSK application, *rab3^rup^*NMJs displayed a slight drop in miniature amplitude and frequency but, strikingly, no change in EPSC amplitudes (**Fig.1b-e**) or in quantal content (0 min: 108 ± 13.3, 20 min: 111 ± 9.6 SEM, n=8 NMJs, p=0.62 paired t-test).

**Figure 1.**
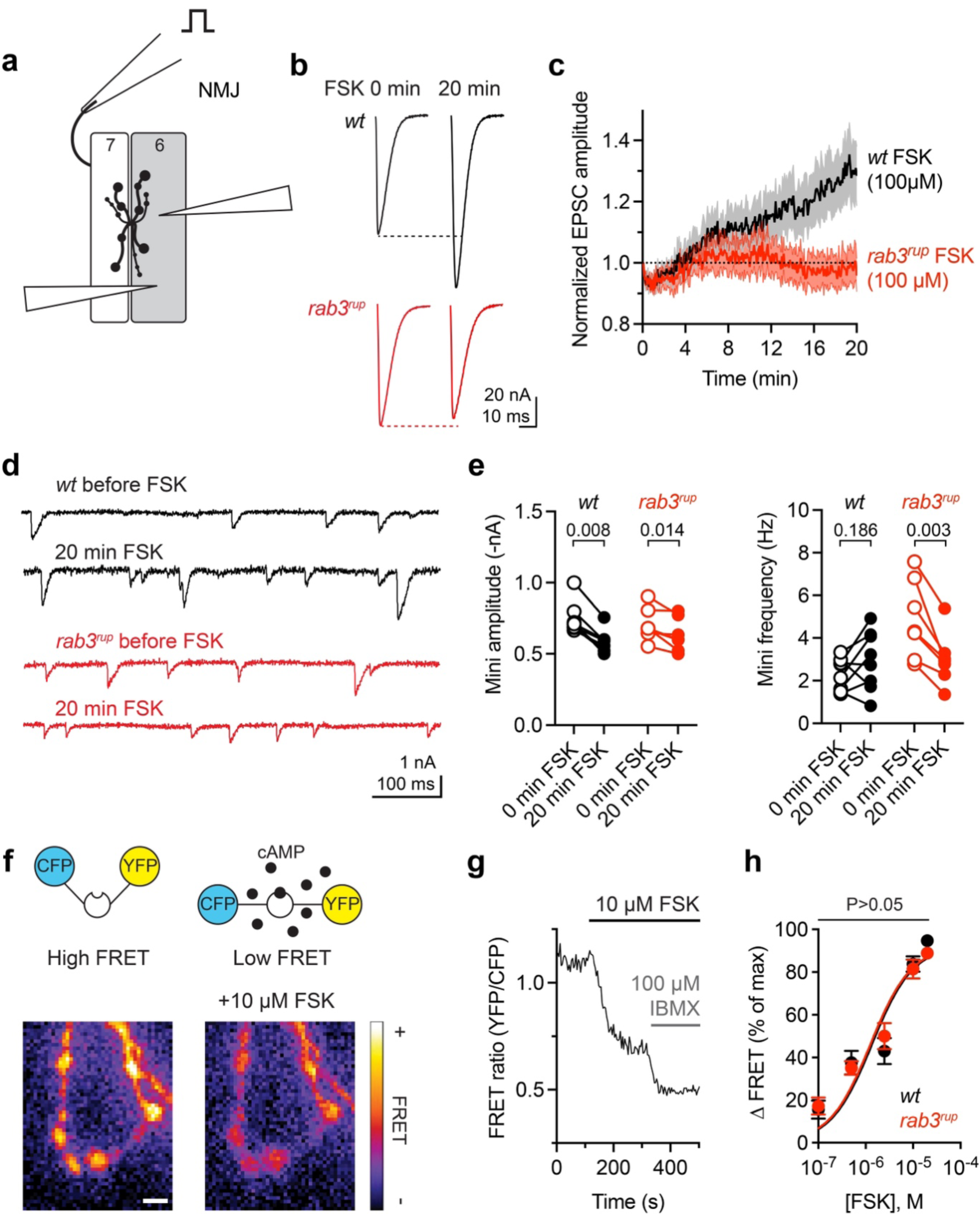
Rab3 is required for cAMP-induced presynaptic potentiation. **a,** Two-electrode voltage-clamp configuration at the NMJ. **b,** Example EPSCs of *wt* (black) and *rab3^rup^*(red) before and 20 min after 100 µM FSK application. **c,** Quantification of EPSC amplitudes. Data (*wt* n=13, *rab3^rup^* n=8 NMJs) are normalized to the initial amplitude and plotted as mean ± SEM. **d,** Example miniature traces of *wt* (black) and *rab3^rup^* (red) before and 20 min after 100 µM FSK application. **e,** Quantification of miniature (mini) amplitudes and frequency (*wt* n=8, *rab3^rup^*n=7 NMJs). P values: Wilcoxon matched-pairs (wt amplitude) or paired t-test (all others). **f,** Epac1-camps scheme and pseudocolour FRET images (YFP/CFP ratios) of motoneuron terminals (*dvglut-GAL4>UAS-Epac1-camps*) with low and high cAMP concentrations (10 µM FSK). Scale bar 2 µm. **g,** Absolute FRET values (YFP/CFP ratios) recorded at an example *wt* NMJ upon addition of FSK and subsequently 100 µM IBMX (3-isobutyl-1-methylxanthin), a non-selective phosphodiesterase inhibitor used to induce a maximal response^60^. **h,** The individual points of the concentration-response curves obtained from traces as in **g** do not differ significantly between *wt* and *rab3^rup^* NMJs (t-test). Data are presented as mean ± SEM.

In principle, these results may be explained by altered cAMP dynamics in *rab3^rup^* mutants. We therefore used the FRET-based cAMP sensor Epac1-camps^13^ to directly measure cAMP production (**Fig. 1f**). FSK application produced similar FRET changes in control and *rab3^rup^*motoneurons, indicating that Rab3 does not influence cAMP levels *per se* (**Fig. 1g,h**).

### Optogenetic cAMP production increases vesicle fusogenicity through Rab3

To improve the spatial precision of cAMP production we used an optogenetic approach by genetically targeting the photoactivatable adenylyl cyclase from the soil bacterium *Beggiatoa* bPAC^wt^ (ref.^14^) to motoneurons (*ok6-GAL4* driver, short: *pre>bPAC^wt^*). This allowed us to elevate cAMP levels with light exclusively in presynaptic cells (**Fig. 2a**). In control animals, continuous photostimulation of bPAC^wt^ for 20 min led to a rapid and sustained >20-fold increase in miniature frequency (**Fig. 2b,c; Extended Data Table**) and rapid, transient potentiation of evoked EPSC amplitudes followed by strong depression (**Fig. 2d**). In contrast, *rab3^rup^*NMJs displayed no change in miniature frequency nor evoked release upon bPAC^wt^ activation (*rab3^rup^, pre>bPAC^wt^*; **Fig. 2b,c,e; Extended Data Table**).

**Figure 2.**
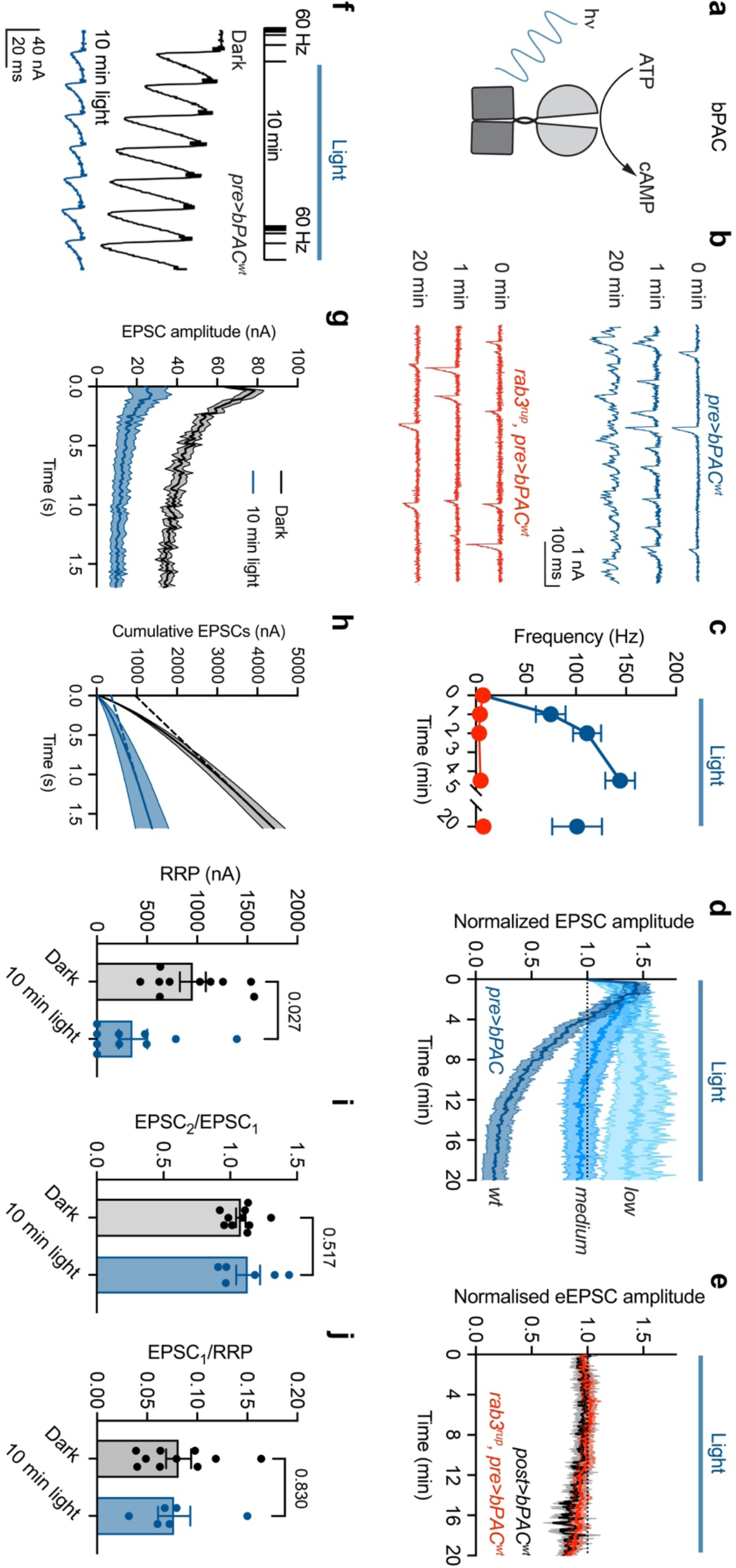
Rab3 mediates a cAMP-dependent enhancement of miniature release and depression of evoked release. **a,** Photo-induced cAMP production by bPAC. **b,** Example taces and **c,** quantification of miniature frequency upon light-tiggered cAMP elevation in *wt* (blue) and *rab3^rup^* (red) motoneurons. **d,** EPSCs (0.2 Hz) normalized to the initial amplitude during 20 min light stimulation of bPAC variants in motoneurons of controls (n=9-10 NMJs). **e,** bPAC^wt^ expressed in the postsynapse (black, n=8) and *rab3^rup^* presynapse (red, n=10). **f,** Train (100 pulses at 60 Hz) and recovery protocol applied twice, separated by 10 min light stimulation. Example traces of the first seven EPSCs with stimulation artefacts removed for clarity. Scale bars 40 nA, 20 ms. **g,** EPSC amplitudes in train (n=10) and **h,** cumulative plot with back-extrapolation to y-axis for RRP estimates before (grey) and after 10 min illumination (blue). **i,** The ratio of the first two EPSCs and **j,** of the first EPSC to the RRP indicate an unaltered p_r_ after 10 min light-induced cAMP production. Data are presented as mean ± SEM. P values (h): Wilcoxon matched-pairs; (i,j): t-test.

We reasoned that the EPSC depression at control NMJs was caused by high photo-induced cAMP concentrations, which exceed those achieved by pharmacological stimulation of endogenous adenylyl cyclases^14,15^. To test this, we utilized two recently engineered bPAC variants with low [bPAC^low^ (R278A): 1.6 ± 0.3 molecules cAMP per molecule PAC per min] and medium [bPAC^medium^ (F198Y): 17 ± 0.8 min^−1^] light-mediated enzymatic activity compared to the wt-version (bPAC^wt^: 93 ± 9 min^−1^; ref.^15^). Consistent with cAMP concentration-dependent presynaptic depression, the light-induced decline of EPSCs correlated with the turnover rate of the cyclases (**Fig. 2d**). In contrast, EPSC amplitudes were unaffected by postsynaptic cAMP production (*g7-GAL4* driver, short: *post>bPAC^wt^*) confirming the second messenger’s presynaptic site of action (**Fig. 2e**).

In order to obtain more mechanistic information on the cAMP-induced changes of evoked release we applied trains of high-frequency stimulation in animals expressing bPAC^wt^ in motoneurons (**Fig. 2f**). After 10 min of light stimulation the amplitude of the first EPSC in the train dropped on average to about 25% of the initial value (**Fig. 2g**). This change was accompanied by a significant decrease in the readily-releasable pool (RRP) of synaptic vesicles, estimated by linear extrapolation of cumulative EPSC amplitudes (**Fig. 2h**)^16^. In contrast, synaptic vesicle release probability (p_r_) appeared unaffected by prolonged cAMP production, as judged by the amplitude of the first EPSC relative to the second EPSC in the train (**Fig. 2i**) and to the total RRP (**Fig. 2j**). This raises the question whether RRP depletion through greatly enhanced miniature neurotransmitter release could explain the gradual depression of evoked release in high cAMP.

To quantitatively test this possibility, we analysed a previously established short-term plasticity model (**Fig. 3a**)^8,17,18^. The model comprises two pools of release-ready vesicles (N_1_ and N_2_) and a pool of supply vesicles (N_0_). N_1_-vesicles have a low p_r_ and recover rapidly, whereas N_2_ vesicles have a high p_r_ and recover slowly^19^. The short-term plasticity model successfully reproduced individual (**Extended Data Fig. 1**) and average high frequency trains with recovery both before and after 10 min of bPAC^wt^ activation (**Fig. 3b**). Consistent with our model-independent analysis (**Fig. 2h-j**), the simulations indicate that cAMP decreases vesicle pool size but has little influence on p_r_ or the rate of vesicle recruitment (**Fig. 3c; Supplementary Table**). We then tested whether miniature release at the experimentally observed frequency (100 Hz; cf. **Fig. 2c**) could deplete the pools of release-ready vesicles and cause depression of evoked release. Keeping all parameters fixed to the values before light stimulation but implementing miniature release from N_2_ at a frequency of only 11 Hz completely exhausted the N_2_ pool and caused a depression comparable to the elevated cAMP condition. The modelling thus demonstrates that even moderate miniature release can cause depression of evoked release when a sub-pool of vesicles possesses a high p_r_ and recovers slowly^19^. However, miniature release from neither N_1_ nor from N_2_, nor from both pools accurately reproduced the short-term plasticity observed upon increased cAMP levels (**Extended Data Fig. 2**). We therefore simulated the cAMP-dependent changes with free model parameters while allowing for RRP depletion by miniatures. Interestingly, the experimental data could be reproduced reasonably well when N_1_ was decreased and the p_r_ of N_1_-vesicles and k_1_ were increased (**Fig. 3d,e; see Methods**). The elevated k_1_ is consistent with a previously reported acceleration of vesicle recruitment by cAMP^20^. Within this framework, the overall p_r_ appears to remain constant despite an increase in p_r_ of the N_1_-vesicles because the N_2_-vesicles with high p_r_ are depleted by miniatures. In summary, the modelling suggests that a cAMP-dependent increase in SV fusogenicity can lead to excessive miniature release and gradual exhaustion of synaptic vesicles available for evoked release with a preferential depletion of high-p_r_ vesicles.

**Figure 3.**
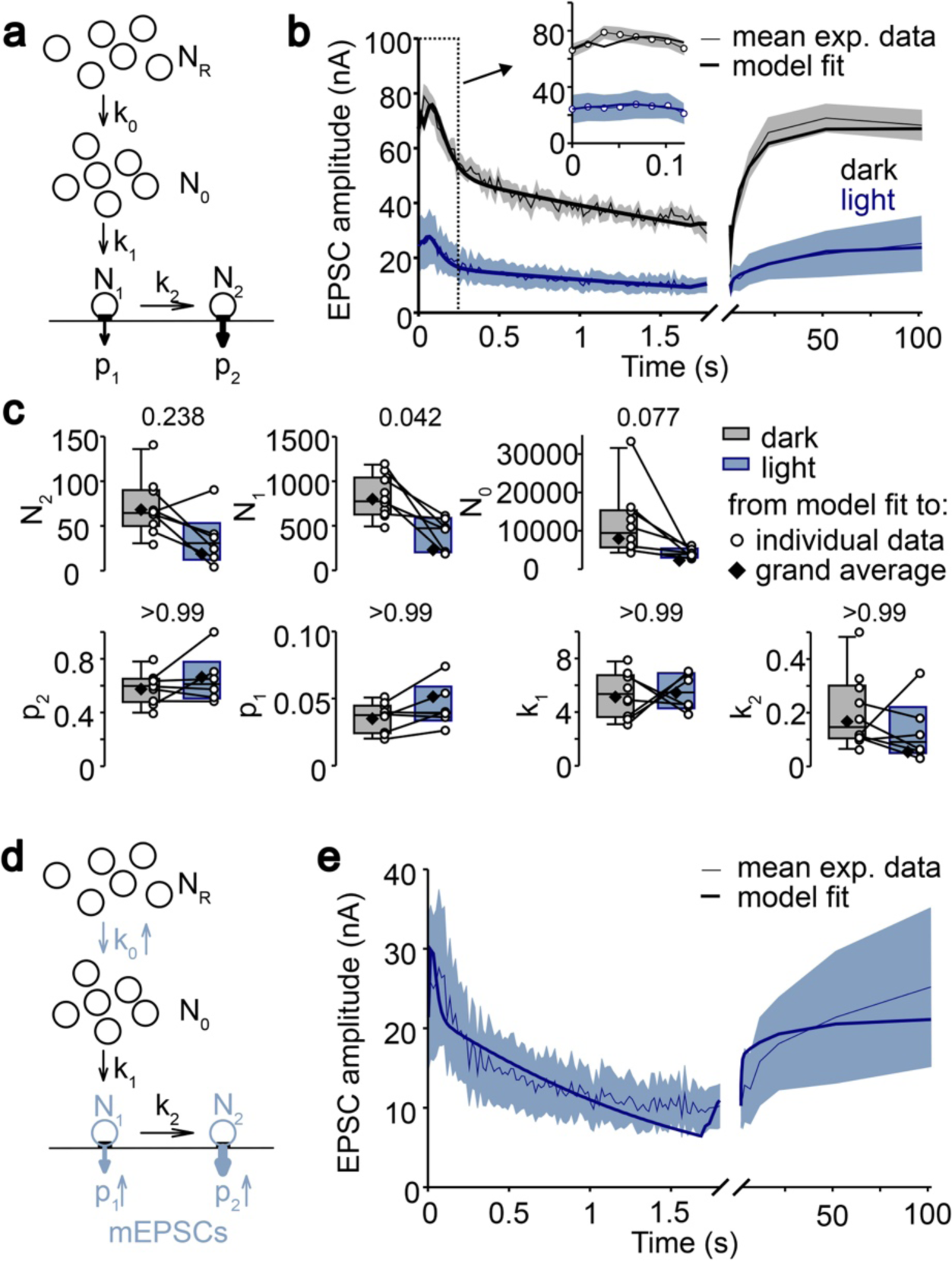
Modelling can explain cAMP-induced synaptic depression. **a,** Schematic illustration of a model containing two fusion-competent vesicle pools with normally primed vesicles (N_1_) exhibiting a low release probability (p_1_) and superprimed vesicles (N_2_) with a high release probability (p_2_), a supply pool (N_0_) and an infinite reserve pool (N_R_). Refilling of N_1_ and N_2_ is determined by the rate constants k_1_ and k_2_, respectively. **b,** Model fit to the mean EPSC train (100 pulses at 60 Hz) and recovery-EPSCs before (grey) and after 10 min illumination (blue). **c,** Individual and median best-fit model parameters before and after 10 min light stimulation. P values: Mann-Whitney U test with a Bonferroni correction (factor 7). **d,** Illustration of altered model parameters and light-induced miniature release from both fusion-competent pools. **e,** Model fit to the mean train of EPSCs (100 pulses at 60 Hz) and recovery-EPSCs after 10 min illumination including increased miniature release. Data are presented as (b,e): model fit (thick line) and mean (thin line) ± SEM (shaded area); (c): box plots (median and quartiles) with whiskers (10-90 percentile) and mean.

### cAMP-induced plasticity correlates with Unc13A repositioning within the active zone

Next, we investigated whether the functional effects of cAMP are accompanied by structural changes in the molecular organisation of AZs. To this end, we employed Stimulated Emission Depletion (STED)^21^ super-resolution microscopy to analyse the nanoscopic arrangement of the scaffolding protein Bruchpilot (Brp)^22^ and the calcium channel subunit Cacophony (Cac)^23,24^ at individual AZs following 20 min of bPAC^wt^ activation (**Fig. 4a**). Neither the AZ area (defined by Brp)^8^, nor the size or number of individual Brp and Cac spots per AZ were altered by high cAMP levels in controls (**Fig. 4b-f**) or *rab3^rup^* animals (**Extended Data Fig. 3; Extended Data Table**). The normal layout of these core components is in line with an intact AZ architecture for neurotransmitter release upon prolonged cAMP production. Aggregates of the AZ protein Unc13 (Munc-13 in mammals) have been suggested to demark synaptic vesicle fusion sites in *Drosophila* and mouse^25,26^. At the *Drosophila* NMJ, synaptic transmission mainly depends on the Unc13A isoform with its exact positioning within the AZ affecting release properties^27,28^.

**Figure 4.**
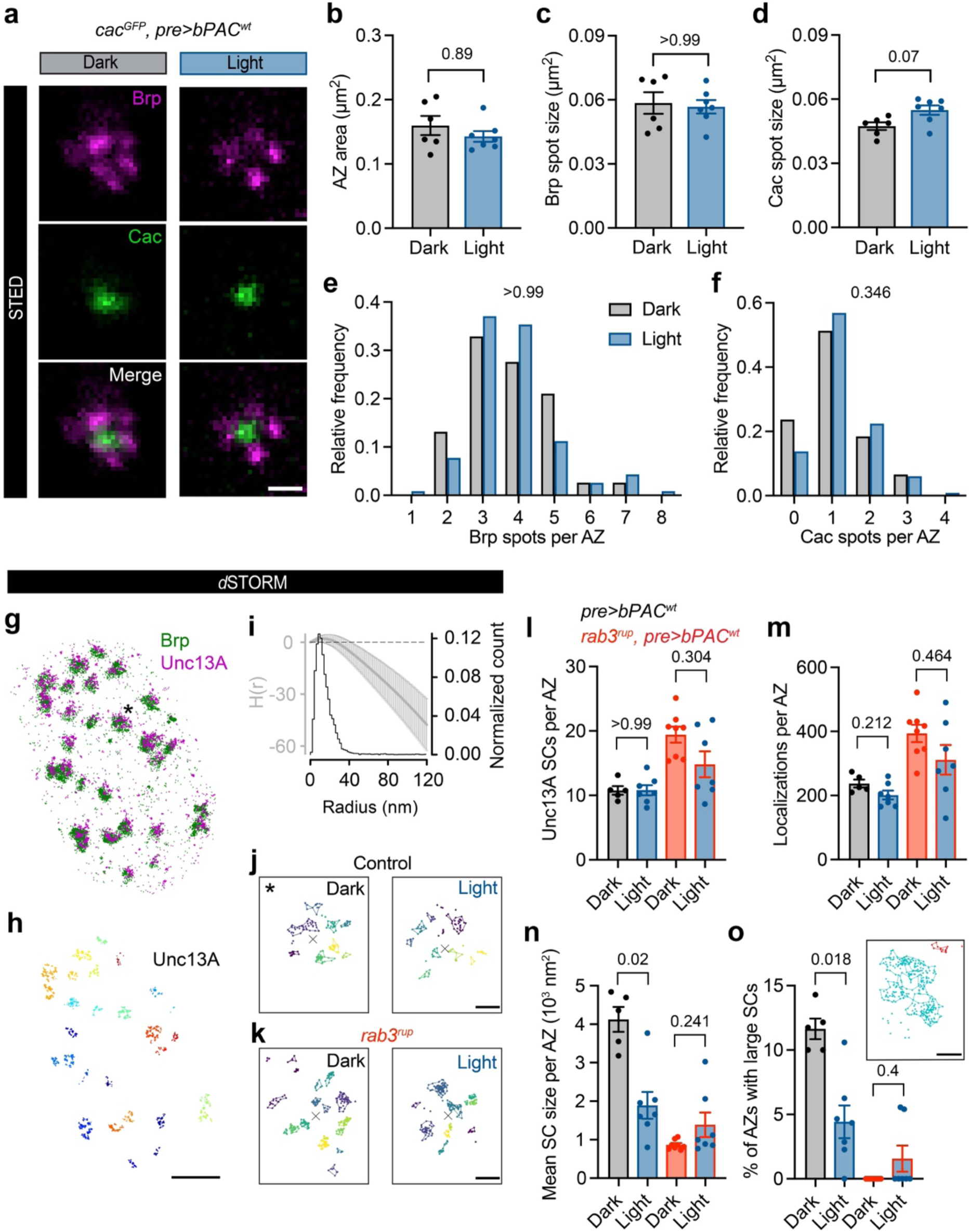
Super-Resolution Microscopy reveals Unc13A reorganization. **a,** Example STED images of Brp (magenta) and Cac (green; staining against GFP-tag of endogenously expressed Cac^24^; *cac^GFP^, pre>bPAC^wt^*) at AZs viewed *en face* (i.e. with the optical axis perpendicular to the AZ membrane) following 20 min light exposure (blue, right) or left in the dark (grey, left). Quantification of **b,** AZ area, **c,** mean spot size of Brp and **d,** Cac (mean ± SEM, dark n=6, light n=7 NMJs), and **e,** spot number per AZ for Brp and **f,** Cac (dark n=76, light n=116 AZs). **g,** Two-channel *d*STORM image of a motoneuron bouton stained against Brp (green) and the Unc13A isoform (magenta). Asterisk marks the enlarged region in (j, dark). **h,** Unc13A localizations with subclusters (SCs) extracted by HDBSCAN and assigned to the nearest Brp-defined AZ (different colours). **i,** Averaged H function (grey, mean ± SD) from n = 503 Unc13A first-level clusters [maximum of the curve indicates a mean SC radius of 13 nm, matching previous work^28^] and histogram (black) of the radius of n = 5805 Unc13A SCs [estimated from SC size assuming a circular area, median (25^th^-75^th^ percentile): 12.3 (8.1-18.3) nm]. Dashed black line, prediction for a random Poisson distribution. **j,** Individual AZ examples showing HDBSCAN-extracted SCs (coloured lines indicate alpha shapes used to determine areas, x marks the AZ centre of mass) with and without light-stimulation in controls and **k,** *rab3^rup^* mutants. **l,** Quantification of average SC number per AZ, **m,** Unc13A localization counts per active zone, **n,** mean SC size per AZ, and **o,** percentage of active zones with large Unc13 clusters (mean size >10.000 nm^2^). Inset: AZ with a small (red) and a large SC (turquoise). Data are presented as mean ± SEM (control: dark n=5, light n=7 NMJs, *rab3^rup^*dark n=8, light n=7 NMJs). Scale bars (a): 250 nm; (g,h): 1 µm; (j,k,o): 100 nm. P values: Mann-Whitney U test with a Bonferroni correction (factor 2).

We therefore employed *direct* Stochastic Optical Reconstruction Microscopy (*d*STORM) to obtain information on the precise spatial organization of Unc13A proteins via the imaging technique’s single-molecule sensitivity^8,29^. In good agreement with previous *d*STORM experiments^28^, single-molecule localization events were arranged in subclusters with a mean radius of 13 nm (**Fig. 4g-k)**. In controls, neither the number of Unc13A subclusters per AZ, nor the number of localizations per AZ, a correlate of protein numbers^8^, were altered by light stimulation (**Fig. l,m**). However, cAMP led to a rearrangement of Unc13A, which resulted in a smaller average subcluster size (**Fig. 4n**). This effect was mediated by a drop in the fraction of AZs containing large Unc13A subclusters (>10,000 nm^2^; **Fig. 4o**). The Unc13 reorganization bears a resemblance to AZ compaction, where an increase in the density of different AZ molecules has been linked to elevated neurotransmitter release^28,30–32^. In contrast, cAMP did not change the area occupied by Unc13A in *rab^rup^* mutants (**Fig. 4n,o**). In summary, we found no evidence for major disruptions of the molecular layout of the AZ or a removal of release sites that would account for the cAMP-induced reduction of evoked transmitter release in control animals. Instead, our results can be explained by increased synaptic vesicle fusogenicity triggered by high cAMP levels, correlating with Unc13A repositioning, and mediated by Rab3. This effect initially potentiates EPSCs before pool depletion sets in through insufficiently constrained spontaneous synaptic vesicle fusions.

### Rab3 is critical for associative learning

Signalling by cAMP plays an important and evolutionarily conserved role in memory formation^2^. In *Drosophila*, odour learning is associated with cAMP-dependent plasticity at presynaptic sites of mushroom body Kenyon Cells (KCs)^33^. Based on our findings at neuromuscular synapses, we therefore asked whether Rab3 is required for olfactory short-term memory. Indeed, aversive learning was abolished in both adult and larval *rab3^rup^* mutants (**Fig. 5a,b; Extended Data Fig. 4**). To narrow down a learning-specific function of Rab3 and distinguish this from brain-wide effects of the null mutant, we used cell-targeted mutagenesis by CRISPR/Cas9 to knock out *rab3* exclusively in ɣ neurons^34^, a sub-population of KCs important for short-term memories^35^ (**Fig. 5c,d**). Disrupting Rab3 function specifically in ɣ KCs (*rab3^CRISPR^*: *GMR71G10-GAL4, 10xUAS-mCD8::GFP/+; UAS-Cas9.C/U6:3-Rab3.gRNA*) significantly impaired odour learning (**Fig. 5e**), further supporting a direct involvement of Rab3 in memory processes. To next test whether the presence of Rab3 only in KCs is sufficient for olfactory learning we re-expressed *rab3* in the mutant background^9^. Driving *UAS-rab3* with a KC-specific *GAL4*-line (*MB247-GAL4*) did not improve the learning scores of *rab^rup^*mutants (**Fig. 5f**). Pan-neuronal re-expression of Rab3 (*elav-GAL4*) gave a partial rescue, illustrating that transgenic expression of *rab3* via the *GAL4/UAS* system can at least partially restore olfactory learning (**Fig. 5g; Extended Data Table**). Taken together, we conclude that Rab3 expression in KCs is necessary but not sufficient for normal learning.

**Figure 5.**
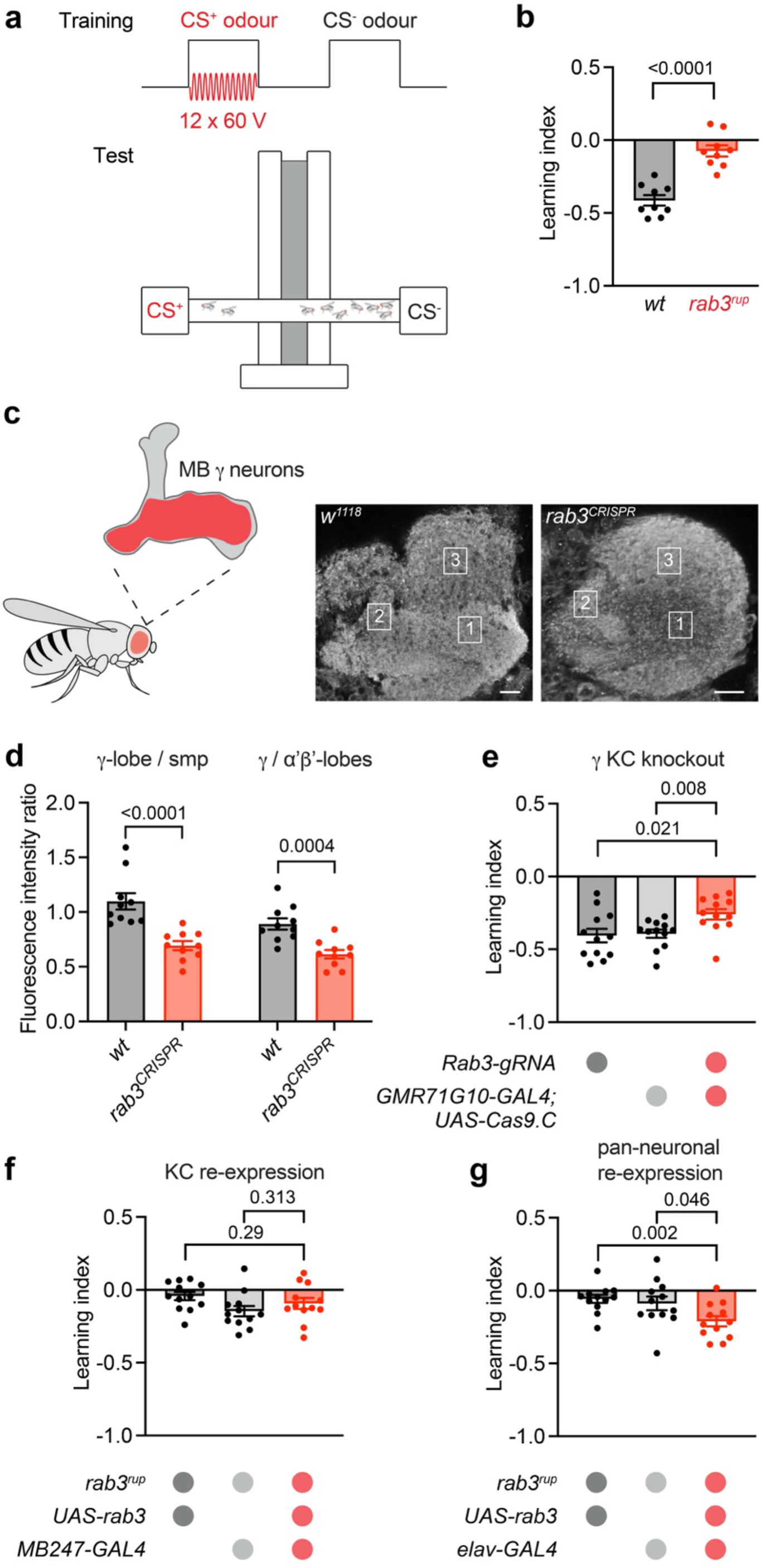
Rab3 is necessary for short-term olfactory learning. **a,** Scheme of the protocol and the apparatus for olfactory classical conditioning using electroshock-conditioned (CS^+^) and unconditioned (CS^−^) odours. **b,** Short-term aversive learning is disrupted in *rab3^rup^* flies (n=9 experiments per genotype). **c,** Illustration of mushroom body (MB) ɣ neurons (red) and antibody staining against Rab3 with ɣ-lobe (1), α‘β’-lobes (2), and superior medial protocerebrum (smp; 3) indicated. **d,** CRISPR/Cas9-mediated knockout of *rab3* specifically in ɣ neurons (red; *rab3^CRISPR^*) decreases the antibody signal in the ɣ-lobe (n=10 brains per genotype) and **e,** significantly reduces learning compared to genetic controls (grey; n=12 experiments per genotype). **f,** Whereas *rab3* re-expression in KCs (*MB247-GAL4*) does not rescue learning in *rab3^rup^*, **g,** panneuronal re-expression (*elav-GAL4*) significantly improves learning scores compared to controls (n=12 experiments per genotype). Scale bars 10 µm. All data are presented as mean ± SEM, P values: t-test.

## Discussion

Here we demonstrate that *Drosophila* Rab3 mediates a cAMP-dependent increase in synaptic vesicle fusion. It was previously shown that Rab3A is important for long-term potentiation at rodent hippocampal mossy fiber synapses, which in turn display a cAMP- and protein kinase A (PKA)-dependent form of presynaptic plasticity^6,36,37^. Unexpectedly, however, Rab3A was found to be dispensable for FSK-induced presynaptic enhancement at this synapse^6^. A possible explanation for this apparent mismatch is that cAMP dynamics during LTP are not mirrored accurately by pharmacological tools like FSK, which may also increase release via Rab3A-independent mechanisms not recruited during mossy fiber LTP^38,39^. In light of the current results, it may therefore be of interest to revisit this topic with the new optogenetic tools now available for spatially and temporally improved control of cAMP production^15,40^. In contrast to the rodent data, *Drosophila* Rab3 is necessary for pharmacologically-induced presynaptic plasticity (**Figure 1b,c**). Thus, our findings provide a coherent model, which causally links Rab3 function to cAMP-induced synaptic plasticity and cAMP-dependent learning and memory. While these results may be partially explained by mechanistically distinct forms of presynaptic plasticity at different synapses or evolutionarily differentiated roles of Rab3 in *Drosophila* and rodents, they clearly motivate further exploring the important but insufficiently understood role of all four mammalian Rab3 isoforms^7^.

Our results show that cAMP-dependent potentiation, depression, and increased miniature release all require Rab3 at the *Drosophila* NMJ. Action potential-evoked EPSC amplitudes were potentiated by cAMP, but with increasing second messenger production evoked neurotransmitter release gradually depressed (**Fig. 2d**). The modelling approach indicates that synaptic depression can be caused by depletion of the available synaptic vesicles through unconstrained miniature release. Indeed, this effect is reminiscent of Rab3A overexpression in PC12 cells, which abolishes Ca^2+^-triggered release by depleting secretory vesicles through constitutive exocytosis^41^. Notably, our interpretation requires at least partial intermixing of vesicle pools for evoked and spontaneous release, which appear to be segregated in other physiological settings^42–44^. Mutants of the SNARE regulator Complexin display a large increase in miniature frequency at the *Drosophila* NMJ without major synaptic vesicle depletion^45,46^. Possibly because the reported frequencies (∼60-80 Hz) are only half as high as those reached through bPAC stimulation or because the N_1_ pool with low p_r_ is primarily affected by spontaneous release in the absence of synaptic vesicle clamping by Complexin.

The involvement of cAMP-mediated synaptic plasticity in learning and memory has been well established^2,47^. In *Drosophila*, associative olfactory conditioning during aversive learning leads to cAMP production in KCs and synaptic depression to output pathways that direct approach, thereby skewing the mushroom body network towards odour-avoidance. Likewise, appetitive learning depresses transmission from KCs to avoidance pathways to drive approach behaviour^48–51^. An intriguing corollary of this model is that increased presynaptic cAMP levels provoke synaptic depression. This aspect has received little attention. The present study demonstrates that high cAMP concentrations can in principle lead to a depression of evoked neurotransmitter release. Future work will have to examine whether e.g. local cAMP signalling domains^52^ trigger a related process at KC AZs during olfactory learning.

The precise positioning of Unc13A within the AZ influences the efficiency of neurotransmitter release^25,27^. Our results show that cAMP triggers nanoscopic reorganizations of Unc13A, with AZs displaying a reduced number of large subclusters while retaining Unc13A protein copies (**Fig. 4l-o**). This effect is reminiscent of AZ compaction during presynaptic homeostatic potentiation (PHP) at the *Drosophila* NMJ, where increased synaptic release is also associated with a decrease in the total Unc13 area per AZ^28^. The altered nanotopology may facilitate release-promoting molecular interactions, e.g. with Syntaxin^26^, or alternatively, the repositioning of Unc13A may reflect the formation of such protein complexes. Most Unc13A subclusters contain eight to twelve (25^th^–75^th^ percentile) fluorophore localizations. Based on previous calculations^8^, it has been estimated that this translates into one or two Unc13 molecules per subcluster^28^. According to the “buttressed-ring hypothesis”, a circular organization of six Munc13 molecules serves as an interaction platform to capture an individual synaptic vesicle and initiate SNARE protein assembly^53^. Thus, the cAMP-induced loss of large subclusters observed in the present study may in fact not reflect compaction, but instead a spreading out of the individual Unc13A molecules. Indeed, this would be predicted upon dilation of the Unc13A-lined vesicular fusion pore followed by collapse of the synaptic vesicle membrane onto the plasma membrane. Large subclusters may therefore demark low-p_r_ presynaptic sites, which are converted to high-p_r_ release sites by cAMP and are infrequently observed at the high-p_r_ AZs of *rab3^rup^* mutants (**Fig. 4o**)^8,9^.

Whereas Rab3 is not a direct phosphorylation target of the cAMP pathway, its interaction partners RIM and Synapsin are PKA substrates^3,54,55^. Synaptic vesicle recruitment from the reserve pool is elevated by cAMP and this process is regulated by Synapsin^20,56^. RIM1 phosphorylation, in turn, has been reported to increase synaptic vesicle docking at the AZ^57^ and to initiate assembly of the tripartite Unc13/RIM/Rab3 complex^58^, which primes synaptic vesicles for fusion in mammals^59^. Both modes of cAMP-dependent presynaptic enhancement likely depend on Rab3 to pass synaptic vesicles on to the secretory machinery. This process may be assisted by Rab3’s suggested role in superpriming^7^. Rab3 is well positioned to mediate presynaptic plasticity due to its dynamic regulation via the GTPase cycle^3^ and its control over the AZ protein composition^9^. The involvement of Rab3 in second messenger signalling by cAMP further underscores its function as an important modulator of neuronal communication.

## METHODS

### Fly stocks

All flies were raised on standard cornmeal and molasses medium at 25°C except for the RNAi knockdown and CRISPR knockout experiments where all genotypes were raised at 29°C two days prior to the experiments. The following fly strains were used:

*w*; ok6-GAL4 w^+^*(ref.^61^)

*w*; g7-GAL4/CyO act-GFP w*^+^

*w^1118^; Df(2R)ED2076, ok6-GAL4 w^+^/CyO GFP w^−^*

*w^1118^; Df(2R)ED2076/CyO GFP w^−^; elav-GAL4*

*w*, dVGlut-GAL4 w^+^; Df(2R)ED2076/CyO GFP*

*w^1118^; UAS-bPAC w^+^/CyO GFP w^−^* (ref.^14^)

*y^1^,w^1118^; 20xUAS-bPAC(R278A)::eYFP/CyO* (RJK559)^15^

*y^1^,w^1118^;; 20xUAS-Venus::bPAC(F198Y)/Sb* (RJK1007)^15^

*w*; rab3^rup^/CyO GFP w^−^; UAS-bPAC/Tb*

*w^1118^;; 20xUAS-epac1(camps) w^+^/Sb* ^13^

*w*; rab3^rup^/CyO GFP w^−^; 20xUAS-epac1(camps)w^+^/Tb*

*w*;; mb247-GAL4 w^+^*(ref.^62^)

*w*; rab3^rup^/Cyo GFP w^−^; UAS-rab3 w^+^* (ref.^9^)

*w*; rab3^rup^/CyO GFP w^−^; MB247-GAL4 w^+^/TM6B,Tb*

*w^1118^, cac^sf-GFP-N^; ok6-Gal4 w^+^/CyO GFP* ^24^

*w^1118^, cac^sf-GFP-N^; Df(2R)ED2076, ok6-Gal4 w^+^/CyO GFP w^−^*

*y,w; GMR71G10-GAL4, 10xUAS-mCD8::GFP/CyO; UAS-Cas9.C/TM6B,Tb* ^34^

BDSC 24635: *w*, dVGlut-GAL4 w^+^;;* ^63^

BDSC 78045: *w*; rab3^rup^* (ref.^9^)

BDSC 458: *elav-GAL4 w^+^;;*

BDSC 81906: *w*;; U6:3-Rab3.gRNA w^+^* (ref.^64^)

VDRC 100787: *w*; UAS-rab3-RNAi/CyO*

### Electrophysiology

Briefly, two electrode voltage clamp (TEVC) recordings were performed on muscle 6 in segments A2 and A3 of wandering third instar male larvae using an Axoclamp 900A amplifier (Molecular devices) with intracellular electrodes of 10-20 MΩ resistance. The measurements were performed at room temperature (RT) in hemolymph-like solution (HL-3)^65^ composed of (in mM): NaCl 70, KCl 5, MgCl_2_ 20, NaHCO_3_ 10, trehalose 5, sucrose 115, HEPES 5, and CaCl_2_ 1.0, pH adjusted to 7.2. Only muscle cells with a membrane potential between −50 and −70 mV and input resistances > 4 MΩ were accepted for analysis. Minis and evoked EPSCs were recorded at −80 mV and −60 mV, respectively. For evoked EPSCs, the innervating nerve was stimulated with 300 µs pulses of 8-15 V (S48/S88 Grass Instruments) via a suction electrode. For cAMP production, bPAC was activated with a blue LED (∼92 µW/mm^2^ at 470 nm; CoolLED or Minostar LED). Signals were sampled at 10 kHz, low-pass filtered at 1kHz, and analysed in Clampfit 10.7 (Molecular Devices). In the FSK experiments (**Fig. 1d**), miniature release was quantified automatically via “template detection” in Clampfit, whereas in the bPAC experiments (**Fig. 2b,c**), miniatures were detected manually in a 500 ms time window to capture the high frequency. Linear fits to EPSCs 80-100 of the cumulatively plotted amplitudes were back extrapolated to estimate RRP sizes in **Fig. 2h**. For EPSC ratio measurements in **Fig. 2i and j**, the amplitude of the second response in the train was measured from the peak to the point of interception with the extrapolated first response. Measurements exhibiting motoneuron recruitment errors or exceeding a holding current of 10 nA were discarded. One NMJ was recorded per animal.

### FRET Imaging

Ratiometric FRET imaging was performed using an upright epifluorescence microscope (BX51WI, Olympus) equipped with a water-immersion objective (60 x, numerical aperture 1), a LED light source (pE-4000, CoolLED), a 445LP dichroic mirror, a beam splitter (Optosplit II, Cairn Research) with a 505LP dichroic mirror and emission filters for CFP (480/30) and YFP (535/30), and an electron-multiplied charge-coupled device (EMCCD) camera (iXon DU-897, Andor). CFP and YFP images upon CFP excitation (435 nm) were captured every 4 s with 80 ms of illumination time. FRET was monitored in real time with the VisiView Software (Visitron) as the ratio between YFP and CFP emissions. The YFP emission was corrected for the spectral bleedthrough of CFP emission into the YFP channel, as previously described^66^. Larvae expressing Epac1-camps in motoneurons were prepared as described for electrophysiology. NMJs on muscles 6/7 in segments A2 and A3 were imaged at RT in HL-3 and stimulated with different FSK concentrations (100 nM, 500 nM, 2.5 µM, 10 µM, 20 µM). Responses to FSK were normalized to the baseline FRET ratio and expressed as percentages of the subsequent response to 100 µM IBMX (20 data points were averaged per ratio).

### STED Microscopy

For STED imaging, dissected male larvae were subjected to 20 min optogenetic activation, as for electrophysiology, or left in the dark for 20 min as controls and then fixed for 20 min in Bouin’s (Roth, 6482). Following blocking for 30 min in PBT (PBS with 0.05% Triton X-100, Sigma Aldrich, 9002-93-1) containing 5% normal goat serum (Sigma Aldrich, G9023) at RT, the preparations were incubated at 4 °C overnight with primary antibodies to stain against GFP of the endogenously-tagged Cac^24^ and Brp: mouse-α-Brp (nc82, 1:100, AB_528108)^67^, rabbit-α-GFP (1:250, Life Technologies, AB_2536526). After two brief and six 10 min washing steps in PBT, the preparations were incubated with secondary antibodies for 2 hours at RT: goat-α-HRP-AlexaFluor488 (1:200, JIR/Dianova, AB_2338965), goat-anti-α-mouse-StarRed (1:200, Abberior, AB_3068620), goat-α-rabbitStarOrange (1:200, Abberior, AB_3068622). After another round of washing, the samples were mounted in Vectashield (Vector Laboratories) and stored at 4°C before STED imaging. Images were acquired with an upright STED microscope (Infinity Line, Abberior Instruments) using an 60x/1.42 NA oil immersion objective and a pulsed 775 nm STED laser to deplete the StarRed and StarOrange dyes. The 2D STED images were acquired with the Imspector software and a pixel size of 30 nm x 30 nm, 5 μs dwell time, and 12 line accumulations. For each set of experiments, all genotypes were stained in the same vial and imaged in one session with identical laser settings to ensure comparability. Image analysis was performed with ImageJ (National Institutes of Health) as previously described^45^ focussing on active zones viewed *en face*.

### Confocal Microscopy

Brains (5 to 8 day old flies) were dissected on ice and fixed in 4% paraformaldehyde for 1 h at RT. After two brief and six 10 min washing steps in 0.3% PBT (PBS with 0.3% Triton X-100, Sigma Aldrich), the samples were incubated in ROTI blocking buffer (1:10 in PBT, Roth) overnight and then with primary antibodies for 48 h at 4°C: rabbit-α-rab3 (1:250; provided by S.J. Sigrist, unpublished) and chicken-α-GFP (1:150, Sigma Aldrich, AB_90890). After a further round of washing, the samples were incubated with secondary antibodies overnight: goat-α-rabbit-StarRed (1:200, Abberior, AB_2833015), chicken-AlexaFluor488 (1:200, Life Technologies, AB_2534096), and goat-α-HRP-AlexaFluor488 (1:200, JIR/Dianova, AB_2338965); washed again, mounted in Vectashield (Vector Laboratories), and stored at 4 °C until imaging. GFP and HRP signals were used to identify the γ-lobes in *rab3^CRISPR^* and *w^1118^* genotypes, respectively. Image stacks of whole-mount brains were acquired with an upright STED microscope in confocal mode (Infinity Line, Abberior Instruments) using an 60x/1.42 NA oil immersion objective. Both genotypes were stained in the same vial and imaged in one session with identical laser settings. Image analysis was carried out with ImageJ (National Institutes of Health). Uniform regions of interest (ROI) were used for signal intensity measurements across all images (maximum projections of 5 optical slices spaced 200 nm). The ratios of mean fluorescence intensity were calculated by dividing the mean intensity of the γ-lobe ROI by the α‘β’ ROI or the superior medial protocerebrum ROI.

### *d*STORM

Larvae were prepared and stained as for STED using the following antibodies: mouse-α-Brp (nc82, 1:250, AB_528108)^67^, guinea pig-α-Unc13A (1:250, provided by S.J. Sigrist)^27^, goat-α-mouse-CF568 (1:500, VWR; AB_10559187), and goat-α-guinea pig-AlexaFluor647 (1.500, Life Technologies; AB_141882). The samples were mounted in photoswitching buffer containing 100 mM mercaptoethylamine, oxygen scavenger system [5% (wt/vol) glucose, 5 U/ml glucose oxidase and 100 U/ml catalase], pH 8.0^8^, and imaged on an inverted microscope (Elyra, Zeiss) equipped with an 100x/1.46 NA oil immersion objective. All imaging was performed under highly inclined and laminated optical (HILO) illumination, 640 nm and 561 nm lasers were used for excitation, and 15,000 frames were recorded with 12 ms exposure time on an electron-multiplying CCD camera (iXON DU-897D, Andor Technology). Images were processed with Zen software (black edition, Zeiss). For calibration, performed twice per measurement day, a 2 min video of pre-mounted MultiSpec beads (Zeiss, 2076-515) was acquired at 50 ms exposure, with both the 640 nm and 561 nm lasers. The channels were aligned using the “Affine” method to account for distortions in the horizontal plane. Single-molecule detection and localisation was performed using an 8 pixel mask with a signal-to-noise ratio of 9 (for AF647) or 6 (for CF568) in the “Peak finder” settings while applying the “Account for overlap” function to localise molecules within a dense environment. Fluorescence spots were localized by fitting to a 2D Gaussian function and localizations were subjected to model-based cross-correlation drift correction. Post-rendering, the two channels were aligned using the affine table values generated during calibration. All preparations were stained in the same vial and image acquisition alternated between control and light-stimulated samples.

Data analysis was performed as previously described^28,30,32^ using custom-written code based on the Python implementation of “Hierarchical Density-Based Spatial Clustering of Applications with Noise” (HDBSCAN)^68^. The “LOCAN” package^69^ was used to load localization tables from Zen software. HDBSCAN parameters “minimum cluster size” and “minimum samples” were 100 and 25 localizations, respectively, for extraction of Brp clusters in the CF568 channel, and 6 and 2 localizations, respectively, for extraction of Unc13A subclusters in the AlexaFluor647 channel. These values were optimized to yield subcluster radii that match the H function (derivative of Ripley’s K function) maximum. Focussing on AZs viewed *en face* (indicated by circularity values ≥0.6), the Brp signal served as a mask to define individual AZs for Unc13A analysis. Denoising of Unc13A localizations was performed based on the Euclidian distance to Brp (≤20 nm)^32^. 2D alpha shapes were used to quantify subcluster areas using the Python version of CGAL (Computational Geometry Algorithms Library; https://www.cgal.org). For alpha shapes of Unc13A subclusters, we chose α-values of 300 nm^2^.

### Modelling

We used a model with two pools of release-ready vesicles and heterogeneous p_r_ similar as described in (ref.^8^) and (ref.^45^). The model consisted of two pools of release-ready vesicles (N_1_ and N_2_) with release probabilities p_1_ and p_2_, respectively, and a supply pool N_0_ (**Fig. 3a**). N_2_ is refilled with rate k_2_ from N_1_, N_1_ is refilled with rate k_1_ from N_0_, and N_0_ is refilled with rate k_0_ from an infinite reserve pool of SVs according to the following differential equations:

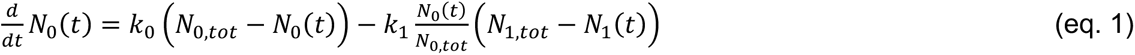

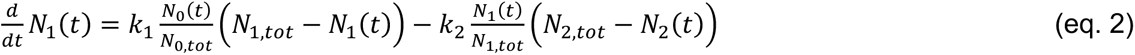

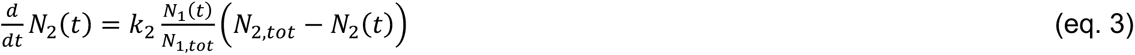

The model also contained a phenomenological description of facilitation as described previously^70^, where each action potential increases both p_2_ and p_1_ by the amount p_x,intial_ (1-p_x_), with x=1 and 2. p_2_ and p_1_ then decay mono-exponentially back to p_x,initial_ with a time constant of τ_f_.

The best-fit parameters (**Fig. 3c**) were determined with a simplex minimization algorithm to reproduce both individual (**Extended Data Fig. 1**) and average EPSC amplitude data of 10 recordings before and 6 recordings after optogenetic cAMP production (**Fig. 3b**). The quality of the fits was calculated by the sum of the squared differences between model fit and the experimental data. Because of the mechanistic importance of the paired-pulse ratio and the time course of recovery from depression, weighting factors were applied when calculating the sum of the squared differences. In particular, the first and the second EPSCs were weighted with the factor of 20 and 5, respectively, and the EPSCs during the recovery from depression were weighted with a factor of 3. The data allowed reliable determination of seven free parameters: N_0_, N_1_, N_2_, k_1_, k_2_, p_1_, and p_2_, i.e., the best-fit parameters were only marginally dependent on the start values. The parameter k_0_ was constrained to 0.031 s^−1^ and τ−_f_ to 0.1 s^45^. For simplicity, the best-fit parameters in **Fig. 3c** for the size of the synaptic vesicle pools (N_x,tot_) were denoted as N_x_, with x = 1 to 3.

To investigate the impact of miniature EPSCs on evoked release, a constant release rate was added either from only N_2_ (**Extended Data Fig. 2a**), from only N_1_ (**Extended Data Fig. 2b**), or from both N_1_ and N_2_ (**Extended Data Fig. 2c**). In these simulations, all other parameters were constrained to the best-fit parameters before light stimulation. Because the miniature release caused the depletion in the initial steady-state condition before evoked release, the initial occupancy was calculated using an analytical solution of the differential equation for the steady-state condition (i.e. setting 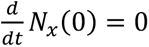, for x = 1 to 3). We assumed an overall miniature release rate upon elevated cAMP levels of 100 s^−1^ (cf. **Fig. 2c**). In the case of miniature release occurring only from N_2_, the miniature release rate from N_2_ was manually adjusted to 11.36 s^−1^ to obtain complete depletion of N_2_ (i.e. the analytically calculated steady-state condition N_2_(0) was 0). In the case of miniature release only from N_1_, the miniature release rate from N_1_ was set to 100 s^−1^. And in the case of miniature release from both N_1_ and N_2_, the miniature release rate from N_1_ and N_2_ were manually adjusted to 11.39 s^−1^ and 100-11.39 = 88.61 s^−1^, respectively, to obtain complete depletion of N_2_ (i.e. the analytically calculated steady-state condition N_2_(0) was 0). Finally, we allowed free optimization of the other parameters of the model to best-fit the average EPSC data after light stimulation (**Fig. 3d and e**) and allowed miniature release with a rate of all together 100 s^−1^ from both N_1_ and N_2_. The corresponding miniature release rates had to be re-adjusted iteratively to 9.5 s^−1^ and 100-9.5 = 91.5 s^−1^, respectively, to obtain complete depletion of N_2_. The resulting best-fit parameters were N_1_ = 107.6 (compared to the control value without light stimulation 798.3), p_1_ = 0.211 (compared to 0.035), k_1_ = 74.69 s^−1^ (compared to 5.13 s^−1^), and k_0_ = 0.016 s^−1^ (compared to 0.031 s^−1^).

The model was implemented in C++ using the compiler of XCode 15 on macOSX 14 (Apple Inc., Cupertino, CA, USA). The required computational time for the minimization of the seven free parameters of the models for the EPSC amplitude before and after light stimulation was less than a minute for all recordings. The results of the minimization were visualized with Mathematica 12 (Wolfram Research, Champaign, IL, USA).

### Learning experiments

Groups of ∼150 flies (5 to 8 days old) were trained for associative olfactory short-term learning essentially as previously described^47^ using a modified learning apparatus to perform four experiments simultaneously. Airflow was adjusted to ∼15 L/min, the relative humidity was set at 80%, and all experiments were performed at 25 °C in complete darkness. The odorants ethyl acetate (EA; Sigma, 141-78-6) or iso-amyl acetate (IAA; Sigma, 123-92-2) diluted in paraffin oil (Sigma, 8012-95-1) at a ratio of 1:100 were used as the conditioned stimulus (CS) and 12 electric shocks of 60 V were applied as the unconditioned stimulus (US). The flies were exposed to one of the odours paired with electric shock reinforcement (CS+) for 1 min and 30 s later the second odour was presented for another minute without the electric shock (CS-). The flies were then moved through the elevator to the T-maze where they were presented with both odours simultaneously and tested for odour preference after 2 min. Reciprocal training was performed by switching the CS+ and CS-odours. The Preference Index (PI) for each experiment was calculated as the number of flies on the CS+ side minus the number of flies on the CS-side, divided by the total number of flies. PI = (#CS+ flies - #CS-flies) / (# total flies). The Learning Index (LI) was calculated by averaging two reciprocal experiments. LI= (PI + PI_reciprocal_) / 2.

Aversive larval learning experiments were carried out at RT on petri dishes freshly prepared the day before the experiments with 1.5% pure Agarose (Roth, 9012-36-6) or 1.5% Agarose with 1.5 M NaCl (Roth, 7647-14-5). Odour cups contained amyl acetate (Sigma, 628-63-7) diluted 1:100 in paraffin oil (Sigma, 8012-95-1) and undiluted 3-octanol (Sigma, 589-98-0). Before starting the experiments, balanced odour preference was confirmed. Training comprised three 5 min cycles on pure agarose with one odour and on salt plates with the other odour. The larvae were then placed in the middle of a salt plate with the two odours on either side and their final position was registered after 5 min^71,72^. Each experiment included thirty larvae, reciprocal training and calculation of PI and LI was performed as for adult learning.

### Statistics

Data were analysed using Prism 9 (GraphPad) or Sigma Plot 13 (Systat). Group means were compared by a two-tailed or paired t-test, unless the assumption of normal sample distribution was violated according to the Shapiro-Wilk test. In this case, a non-parametric Mann-Whitney U test or Wilcoxon matched-pairs test was employed. For comparison of the 7 best-fit parameters of the model (**Fig. 3c**) and between dark and light-stimulated STED and *d*STORM images (**Fig. 4b-f, l-o; Extended Data Fig. 3a-e**) a Mann-Whitney U test with a Bonferroni correction was used with a factor of 7 and 2, respectively. Data and statistics are summarized in the Extended Data Table and Supplementary Table.

## ACKNOWLEDGEMENTS

We thank N. Naumann and B. Goettgens for technical support, N. Scholz and A.-K. Dahse for genotyping, S.J. Sigrist for antibodies, and the Bloomington Stock Centre, the VDRC, and O. Schuldiner for fly strains. This work was supported by the Deutsche Forschungsgemeinschaft (KI1460/7-1/SPP 2205, KI1460/9-1/KFO 5001, KI1460/5-1, and INST 268/437-1 FUGG to RJK; FOR 3004 SYNAPBS, HA6386/10-2 to SH; PA1979/3-1 and PA1979/5-1 to DP), the Leipzig University Clinician Scientist Program and the Jung Foundation for Science and Research to AM (Jung Career Advancement Prize 2023), the Nottingham Research Anne McLaren fellowship to IM, and by the European Research Council (ERC Consolidator Grant 865634 to SH).

## AUTHOR CONTRIBUTIONS

D.S: writing–review and editing, investigation, formal analysis, and visualization. A.A.: writing– review and editing, investigation, formal analysis, and visualization. A.M.: writing–review and editing, formal analysis, and visualization. J.N.: writing–review and editing, investigation, formal analysis, and visualization. M.L.: writing–review and editing, investigation. N.H.: writing– review and editing, investigation. N.E.: writing–review and editing, formal analysis. D.P.: writing–review and editing, formal analysis. T.S.: writing–review and editing, investigation, formal analysis, and visualization. I.M.: writing–review and editing, formal analysis. M.S.: writing–review and editing, formal analysis. M.H.: writing–review and editing, formal analysis. S.H.: writing–review and editing, conceptualization, investigation, formal analysis, visualization, and supervision. R.J.K.: writing–original draft, writing–review and editing, initiated the project, conceptualization, formal analysis, visualization, and supervision.

## COMPETING NINTERESTS

The authors declare no competing interests.

## EXTENDED DATA

**Extended Data Figure 1.**
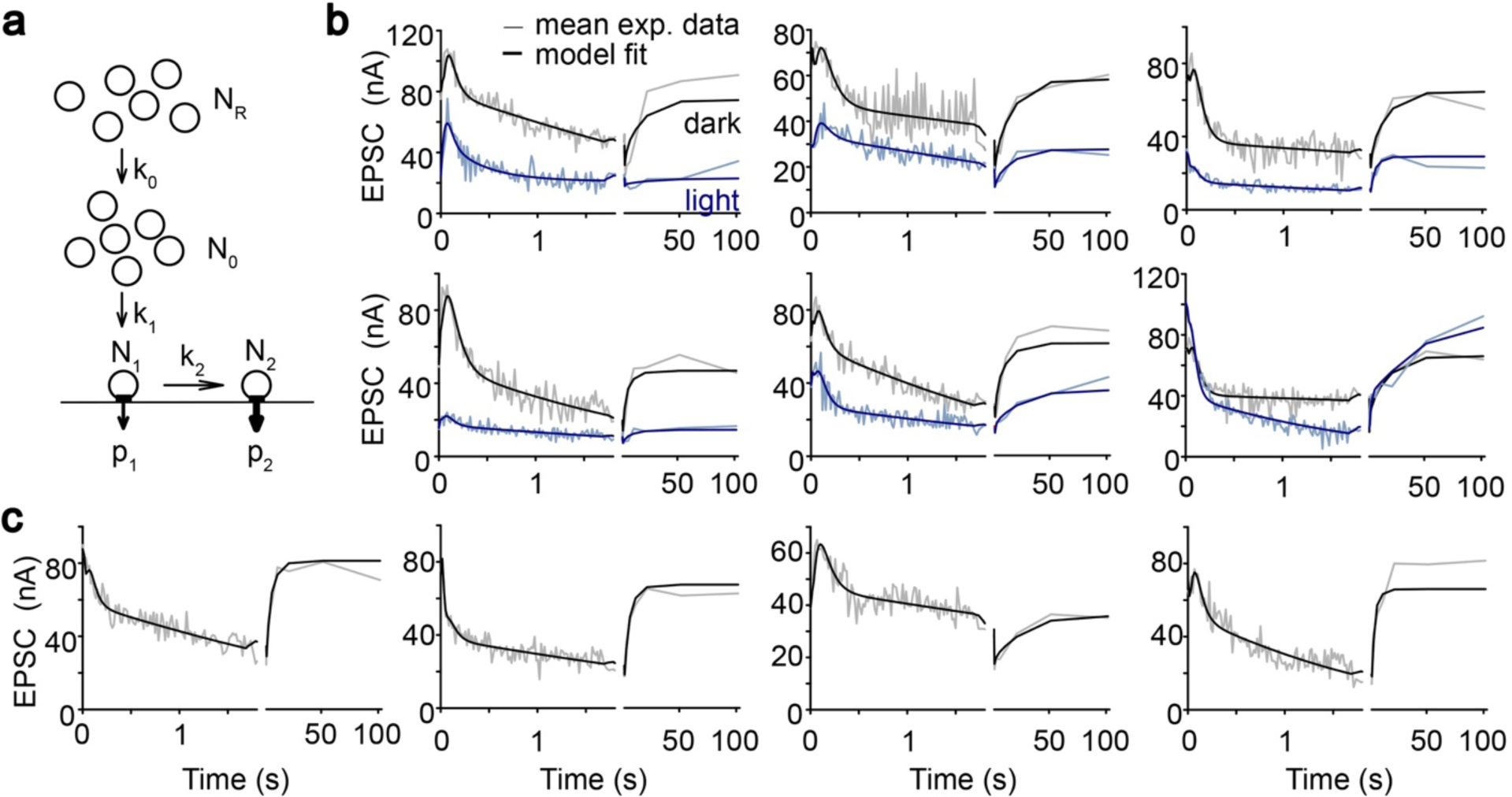
Modelling: individual traces. **a,** Schematic illustration of the model. **b,** Model fits to individual trains of EPSCs (100 pulses at 60 Hz) and recovery-EPSCs before (grey) and after 10 min illumination (blue). **c,** Same as panel (b) for experiments in which only the data before illumination were available. Note that in these cells evoked synaptic transmission was completely abolished after 10 min light stimulation.

**Extended Data Figure 2.**
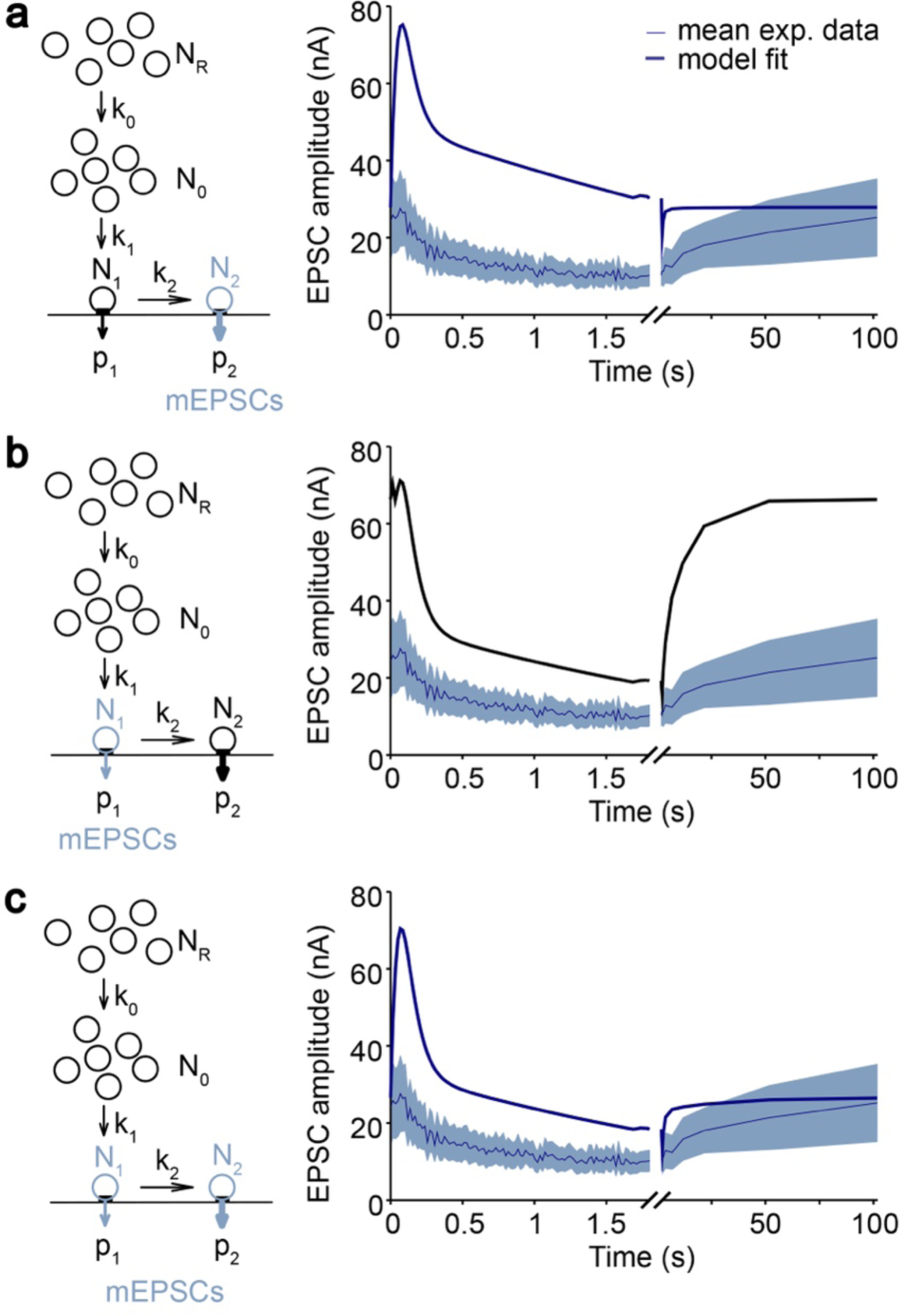
Modelling of miniature release from fusion-competent synaptic vesicle pools. **a,** *Left:* Schematic illustration of the model with miniature release exclusively from the superprimed vesicle pool N_2_. *Right:* Model fit to the mean train of EPSCs (100 pulses at 60 Hz) and recovery-EPSCs after 10 min illumination. **b,** Same as in panel a but for miniature release exclusively from the normally-primed vesicle pool N_1_. **c,** Same as in panel a but for miniature release from both N_1_ and N_2_. Data are presented as model fit (thick line) and mean (thin line) ± SEM (shaded area).

**Extended Data Figure 3.**
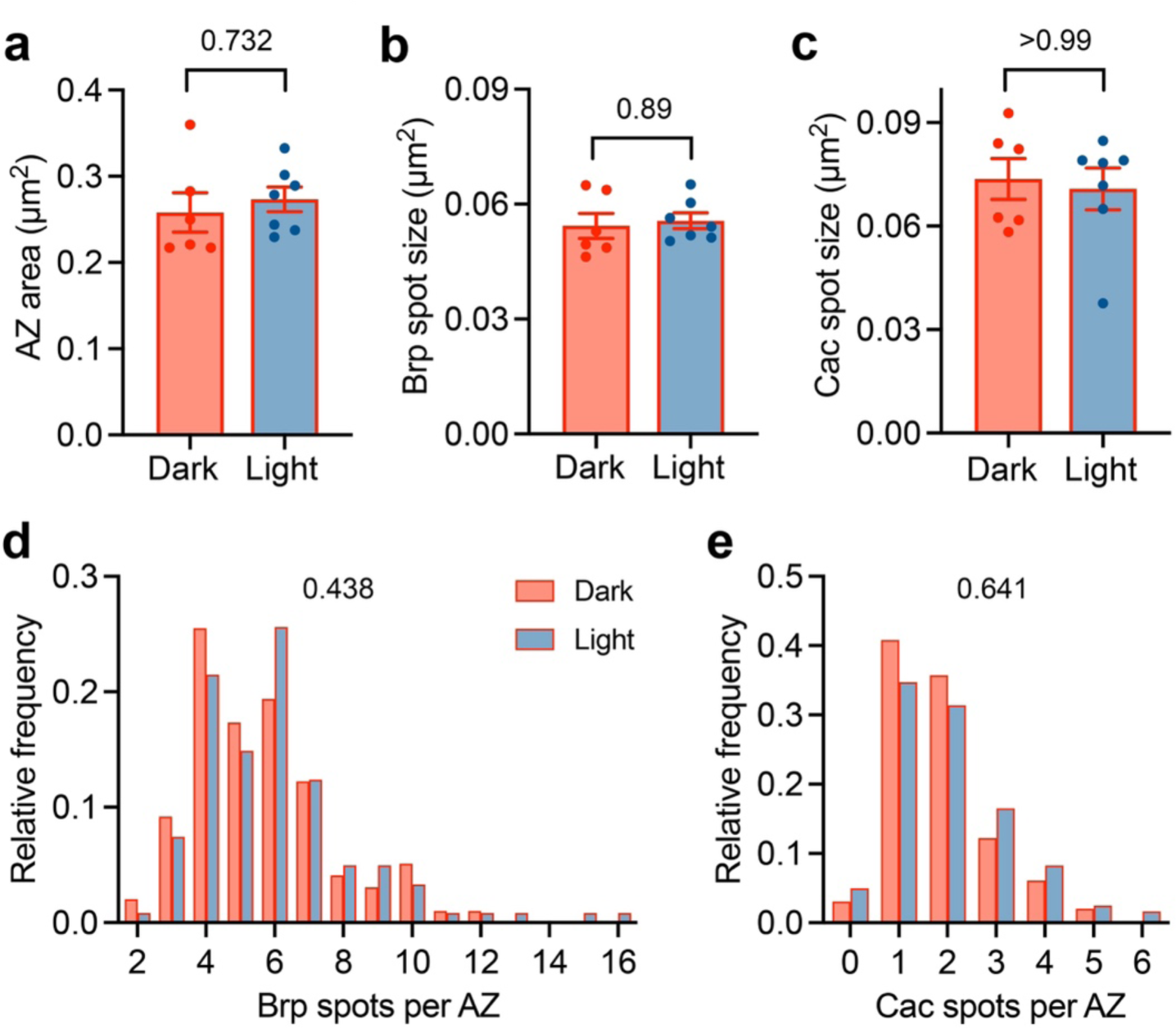
STED Microscopy of *rab3^rup^* AZs. Quantification of **a,** AZ area, **b,** mean spot size of Brp and **c,** Cac (mean ± SEM, dark n=6, light n=7 NMJs), and **d,** spot number per AZ for Brp and **e,** Cac (dark n=98, light n=121 AZs). Genotype: *rab3^rup^, cac^GFP^, pre>bPAC^wt^*. P values: Mann-Whitney U test with a Bonferroni correction (factor 2).

**Extended Data Figure 4.**
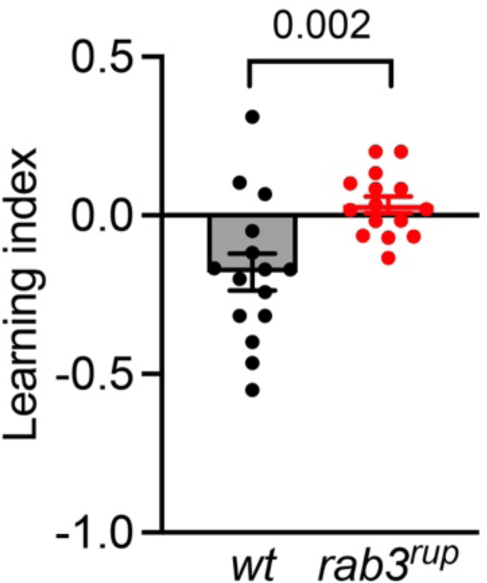
Aversive learning in *Drosophila* larvae. Salt learning is lost in *rab3^rup^* mutant larvae. Data are presented as mean ± SEM, P value: t-test.

**Extended Data Table.**
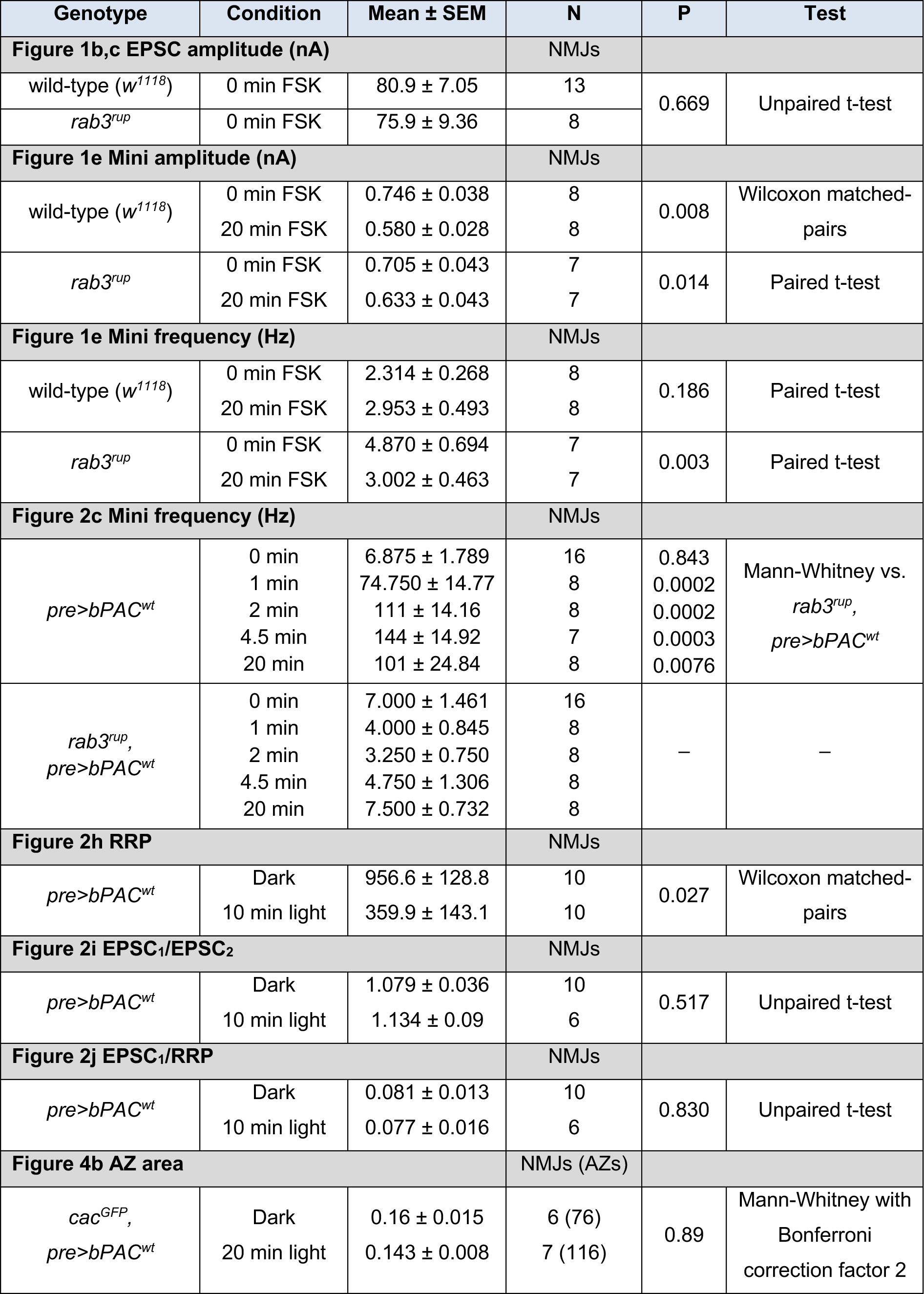
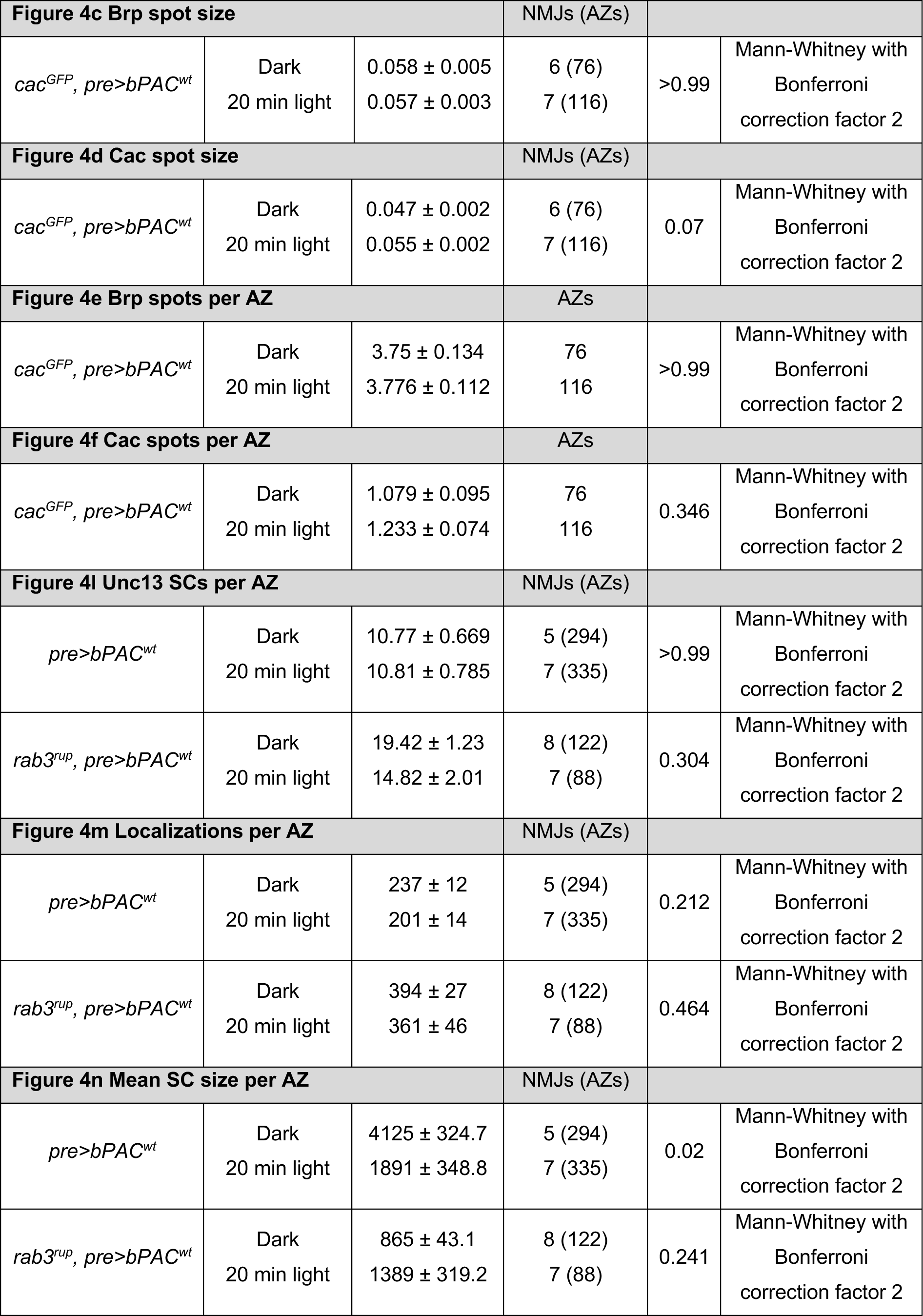
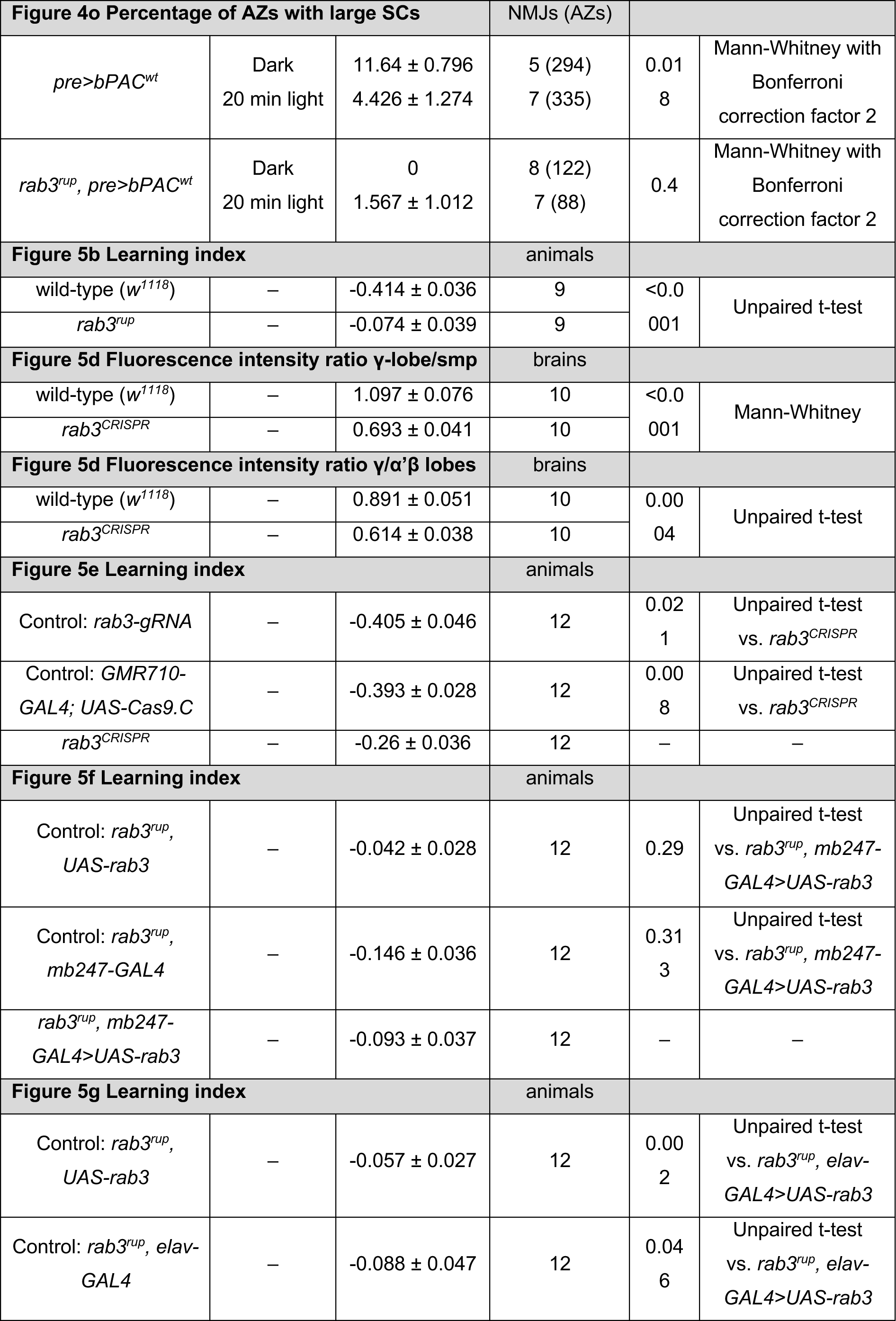
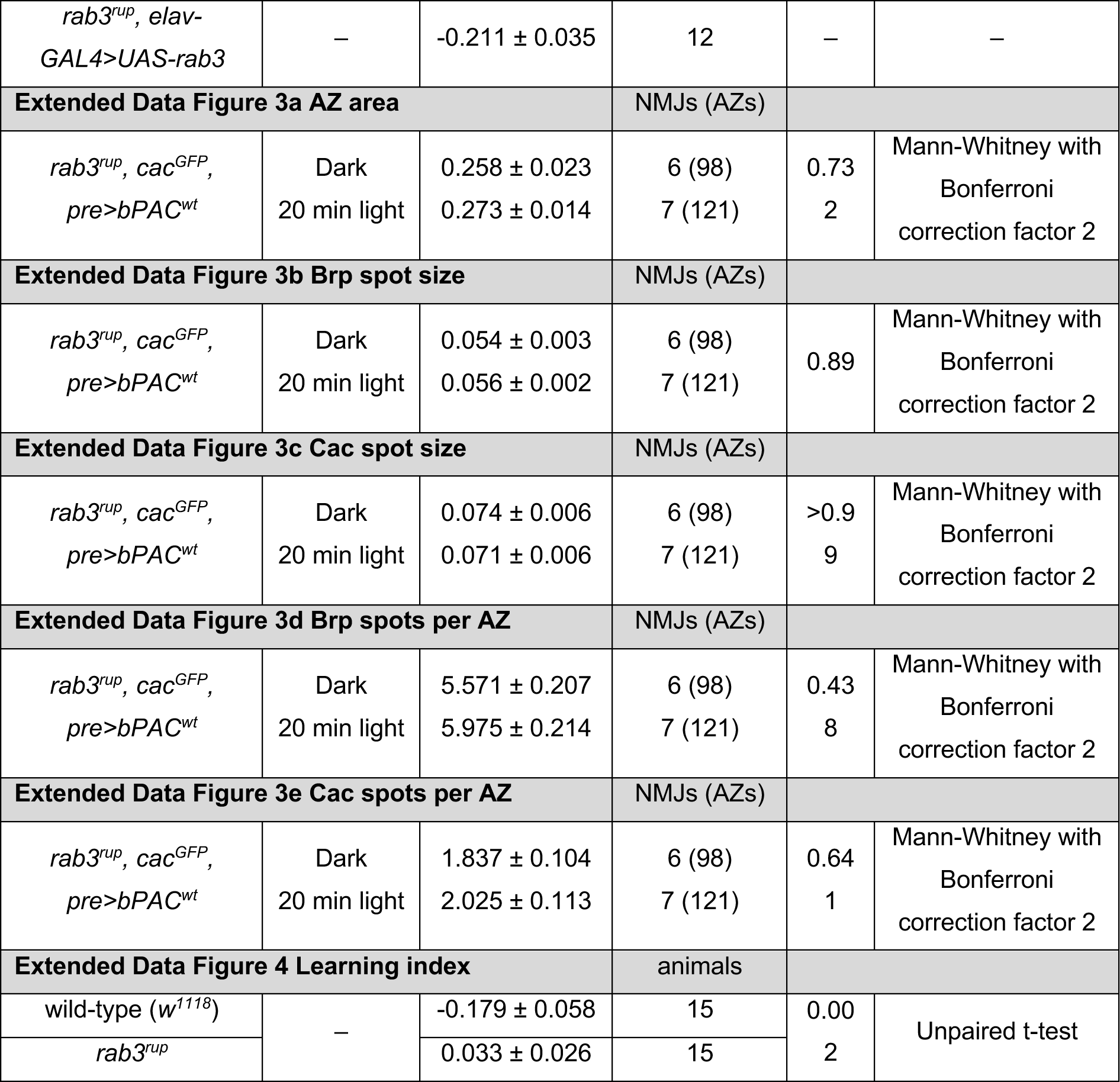

## REFERENCES

1. Monday, H. R., Younts, T. J. & Castillo, P. E. Long-term plasticity of neurotransmitter release: Emerging mechanisms and contributions to brain function and disease. Annu. Rev. Neurosci. 41, 299–322 (2018).

2. Kandel, E. R., Dudai, Y. & Mayford, M. R. The molecular and systems biology of memory. Cell 157, 163–186 (2014).

3. Südhof, T. C. The synaptic vesicle cycle. Annu. Rev. Neurosci. 27, 509–547 (2004).

4. Takamori, S. et al. Molecular anatomy of a trafficking organelle. Cell 127, 831–846 (2006).

5. Tsetsenis, T. et al. Rab3B protein is required for long-term depression of hippocampal inhibitory synapses and for normal reversal learning. Proc. Natl. Acad. Sci. U. S. A. 108, 14300–14305 (2011).

6. Castillo, P. E. et al. Rab3A is essential for mossy fibre long-term potentiation in the hippocampus. Nature 388, 590–593 (1997).

7. Schlüter, O. M., Basu, J., Südhof, T. C. & Rosenmund, C. Rab3 superprimes synaptic vesicles for release: Implications for short-term synaptic plasticity. J. Neurosci. 26, 1239–1246 (2006).

8. Ehmann, N. et al. Quantitative super-resolution imaging of Bruchpilot distinguishes active zone states. Nat. Commun. 5, 4650 (2014).

9. Graf, E. R., Daniels, R. W., Burgess, R. W., Schwarz, T. L. & DiAntonio, A. Rab3 dynamically controls protein composition at active zones. Neuron 64, 663–677 (2009).

10. Leenders, A. G. M., Lopes da Silva, F. H., Ghijsen, W. E. J. M. & Verhage, M. Rab3A is involved in transport of synaptic vesicles to the active zone in mouse brain nerve terminals. Mol. Biol. Cell 12, 3095–3102 (2001).

11. Nonet, M. L. et al. Caenorhabditis elegans rab-3 mutant synapses exhibit impaired function and are partially depleted of vesicles. J. Neurosci. 17, 8061–8073 (1997).

12. Cheung, U., Atwood, H. L. & Zucker, R. S. Presynaptic effectors contributing to cAMP-induced synaptic potentiation in Drosophila. J. Neurobiol. 66, 273–280 (2006).

13. Maiellaro, I., Lohse, M. J., Kittel, R. J. & Calebiro, D. cAMP signals in Drosophila motor neurons are confined to single synaptic boutons. Cell Rep. 17, 1238–1246 (2016).

14. Stierl, M. et al. Light modulation of cellular cAMP by a small bacterial photoactivated adenylyl cyclase, bPAC, of the soil bacterium Beggiatoa. J. Biol. Chem. 286, 1181–1188 (2011).

15. Yang, S. et al. PACmn for improved optogenetic control of intracellular cAMP. BMC Biol. 19, 1–17 (2021).

16. Schneggenburger, R., Meyer, A. C. & Neher, E. Released fraction and total size of a pool of immediately available transmitter quanta at a calyx synapse. Neuron 23, 399–409 (1999).

17. Hallermann, S., Heckmann, M. & Kittel, R. J. Mechanisms of short-term plasticity at neuromuscular active zones of Drosophila. HFSP J. 4, 72–84 (2010).

18. Weichard, I. et al. Fully-primed slowly-recovering vesicles mediate presynaptic LTP at neocortical neurons. Proc. Natl. Acad. Sci. 120, 2017 (2023).

19. Sakaba, T. & Neher, E. Calmodulin mediates rapid recruitment of fast-releasing synaptic vesicles at a calyx-type synapse. Neuron 32, 1119–1131 (2001).

20. Kuromi, H. & Kidokoro, Y. Tetanic stimulation recruits vesicles from reserve pool via a cAMP-mediated process in Drosophila synapses. Neuron 27, 133–143 (2000).

21. Hell, S. W. Far-Field Optical Nanoscopy. Science 316, 1153–1158 (2007).

22. Kittel, R. J. et al. Bruchpilot promotes active zone assembly, Ca2+ channel clustering, and vesicle release. Science 312, 1051–1054 (2006).

23. Kawasaki, F., Felling, R. & Ordway, R. W. A temperature-sensitive paralytic mutant defines a primary synaptic calcium channel in Drosophila. J. Neurosci. 20, 4885–4889 (2000).

24. Gratz, S. J. et al. Endogenous tagging reveals differential regulation of Ca2+ channels at single active zones during presynaptic homeostatic potentiation and depression. J. Neurosci. 39, 2416–2429 (2019).

25. Reddy-Alla, S. et al. Stable Positioning of Unc13 Restricts Synaptic Vesicle Fusion to Defined Release Sites to Promote Synchronous Neurotransmission. Neuron 1350–1364 (2017).

26. Sakamoto, H. et al. Synaptic weight set by Munc13-1 supramolecular assemblies. Nat. Neurosci. 21, 41–55 (2018).

27. Böhme, M. A. et al. Active zone scaffolds differentially accumulate Unc13 isoforms to tune Ca(2+) channel-vesicle coupling. Nat. Neurosci. 19, 1311–20 (2016).

28. Dannhäuser, S. et al. Endogenous tagging of Unc-13 reveals nanoscale reorganization at active zones during presynaptic homeostatic potentiation. Front. Cell. Neurosci. 16, 1–15 (2022).

29. Heilemann, M., et al. Subdiffraction-resolution fluorescence imaging with conventional fluorescent probes. Angew. Chemie - Int. Ed. 47, 6172–6176 (2008).

30. Mrestani, A. et al. Active zone compaction correlates with presynaptic homeostatic potentiation. Cell Rep. 37, (2021).

31. Ghelani, T. et al. Interactive nanocluster compaction of the ELKS scaffold and Cacophony Ca2+ channels drives sustained active zone potentiation. Sci. Adv. 9, eade7804 (2023).

32. Mrestani, A. et al. Nanoscaled RIM clustering at presynaptic active zones revealed by endogenous tagging. Life Sci. alliance 6, 1–14 (2023).

33. Heisenberg, M. Mushroom body memoir: from maps to models. Nat. Rev. Neurosci. 4, 266–275 (2003).

34. Meltzer, H. et al. Tissue-specific (ts)CRISPR as an efficient strategy for in vivo screening in Drosophila. Nat. Commun. 10, (2019).

35. Zars, T., Fischer, M., Schulz, R. & Heisenberg, M. Localization of a short-term memory in Drosophila. Science 288, 672–675 (2000).

36. Weisskopf, M. G., Castillo, P. E., Zalutsky, R. A. & Nicoll, R. A. Mediation of hippocampal mossy fiber long-term potentiation by cyclic AMP. Science 265, 1878–1882 (1994).

37. Huang, Y. Y., Li, X. C. & Kandel, E. R. cAMP contributes to mossy fiber LTP by initiating both a covalently mediated early phase and macromolecular synthesis-dependent late phase. Cell 79, 69–79 (1994).

38. Laurenza, A., Sutkowski, E. M. H. & Seamon, K. B. Forskolin: a specific stimulator of adenylyl cyclase or a diterpene with multiple sites of action? Trends Pharmacol. Sci. 10, 442–447 (1989).

39. Lonart, G., Janz, R., Johnson, K. M. & Südhof, T. C. Mechanism of action of rab3A in mossy fiber LTP. Neuron 21, 1141–1150 (1998).

40. Oldani, S. et al. SynaptoPAC, an optogenetic tool for induction of presynaptic plasticity. J. Neurochem. 156, 324–336 (2021).

41. Schlüter, O. M., Khvotchev, M., Jahn, R. & Südhof, T. C. Localization versus function of Rab3 proteins: Evidence for a common regulatory role in controlling fusion. J. Biol. Chem. 277, 40919–40929 (2002).

42. Crawford, D. C. & Kavalali, E. T. Molecular underpinnings of synaptic vesicle pool heterogeneity. Traffic 16, 338–364 (2015).

43. Melom, J. E., Akbergenova, Y., Gavornik, J. P. & Littleton, J. T. Spontaneous and evoked release are independently regulated at individual active zones. J. Neurosci. 33, 17253–17263 (2013).

44. Peled, E. S., Newman, Z. L. & Isacoff, E. Y. Evoked and spontaneous transmission favored by distinct sets of synapses. Curr. Biol. 24, 484–493 (2014).

45. Scholz, N. et al. Complexin cooperates with Bruchpilot to tether synaptic vesicles to the active zone cytomatrix. J. Cell Biol. 218, 1011–1026 (2019).

46. Huntwork, S. & Littleton, J. T. A complexin fusion clamp regulates spontaneous neurotransmitter release and synaptic growth. Nat. Neurosci. 10, 1235–1237 (2007).

47. Tully, T. & Quinn, W. G. Classical conditioning and retention in normal and mutant Drosophila melanogaster. J. Comp.Physiol. A 157, 263–277 (1985).

48. Owald, D. & Waddell, S. Olfactory learning skews mushroom body output pathways to steer behavioral choice in Drosophila. Curr. Opin. Neurobiol. 35, 178–184 (2015).

49. Handler, A. et al. Distinct dopamine receptor pathways underlie the temporal sensitivity of associative learning. Cell 178, 60–75.e19 (2019).

50. Tomchik, S. M. & Davis, R. L. Dynamics of learning-related cAMP signaling and stimulus integration in the Drosophila olfactory pathway. Neuron 64, 510–521 (2009).

51. Hige, T., Aso, Y., Modi, M. N., Rubin, G. M. & Turner, G. C. Heterosynaptic plasticity underlies aversive olfactory learning in Drosophila. Neuron 88, 985–998 (2015).

52. Anton, S. E. et al. Receptor-associated independent cAMP nanodomains mediate spatiotemporal specificity of GPCR signaling. Cell 185, 1130–1142.e11 (2022).

53. Rothman, J. E., Krishnakumar, S. S., Grushin, K. & Pincet, F. Hypothesis – buttressed rings assemble, clamp, and release SNAREpins for synaptic transmission. FEBS Lett. 591, 3459–3480 (2017).

54. Giovedì, S., Darchen, F., Valtorta, F., Greengard, P. & Benfenati, F. Synapsin is a novel Rab3 effector protein on small synaptic vesicles: II. Functional effects of the Rab3A-synapsin I interaction. J. Biol. Chem. 279, 43769–43779 (2004).

55. Lonart, G. et al. Phosphorylation of RIM1α by PKA triggers presynaptic long-term potentiation at cerebellar parallel fiber synapses. Cell 115, 49–60 (2003).

56. Chi, P., Greengard, P. & Ryan, T. A. Synaptic vesicle mobilization is regulated by distinct synapsin I phosphorylation pathways at different frequencies. Neuron 38, 69–78 (2003).

57. Müller, J. A. et al. A presynaptic phosphosignaling hub for lasting homeostatic plasticity. Cell Rep. 39, (2022).

58. Wang, X. T. et al. Camp−epac−pkcε−rim1α signaling regulates presynaptic long-term potentiation and motor learning. Elife 12, 1–29 (2023).

59. Dulubova, I. et al. A Munc13/RIM/Rab3 tripartite complex: From priming to plasticity? EMBO J. 24, 2839–2850 (2005).

## Methods references

60. Scholz, N. et al. Mechano-dependent signaling by latrophilin/CIRL quenches cAMP in proprioceptive neurons. Elife 6, 1–21 (2017).

61. Sanyal, S. Genomic mapping and expression patterns of C380, OK6 and D42 enhancer trap lines in the larval nervous system of Drosophila. Gene Expr. Patterns 9, 371–380 (2009).

62. Schulz, R. A., Chromey, C., Lu, M. F., Zhao, B. & Olson, E. N. Expression of the D-MEF2 transcription in the Drosophila brain suggests a role in neuronal cell differentiation. Oncogene 12, 1827–1831 (1996).

63. Daniels, R. W., Gelfand, M. V., Collins, C. A. & DiAntonio, A. Visualizing glutamatergic cell bodies and synapses in Drosophila larval and adult CNS. J. Comp. Neurol. 508, 131–152 (2008).

64. Poe, A. R. et al. Robust CRISPR/CAS9-mediated tissue-specific mutagenesis reveals gene redundancy and perdurance in drosophila. Genetics 211, 459–472 (2019).

65. Stewart, B. A., Atwood, H. L., Renger, J. J., Wang, J. & Wu, C. F. Improved stability of Drosophila larval neuromuscular preparations in haemolymph-like physiological solutions. J. Comp. Physiol. A 175, 179–191 (1994).

66. Börner, S. et al. FRET measurements of intracellular cAMP concentrations and cAMP analog permeability in intact cells. Nat. Protoc. 6, 427–438 (2011).

67. Wagh, D. A. et al. Bruchpilot, a protein with homology to ELKS/CAST, is required for structural integrity and function of synaptic active zones in Drosophila. Neuron 49, 833–844 (2006).

68. McInnes, L., Healy, J. & Astels, S. hdbscan: Hierarchical density based clustering. J. Open Source Softw. 2, 205 (2017).

69. Doose, S. LOCAN: A python library for analyzing single-molecule localization microscopy data. Bioinformatics 38, 2670–2672 (2022).

70. Markram, H., Wang, Y. & Tsodyks, M. Differential signaling via the same axon of neocortical pyramidal neurons. Proc. Natl. Acad. Sci. U. S. A. 95, 5323–5328 (1998).

71. Aceves-Piña, E. O. & Quinn, W. G. Learning in normal and mutant Drosophila larvae. Science (80-.). 206, 93–96 (1979).

72. Widmann, A. et al. Genetic Dissection of Aversive Associative Olfactory Learning and Memory in Drosophila Larvae. PLoS Genet. 12, 1–32 (2016).

