## Supplementary Table for "Rab3 mediates cyclic AMP-dependent presynaptic plasticity and olfactory learning"

| | | parameter | | | | | | | | | | | | | |
|---|---|---|---|---|---|---|---|---|---|---|---|---|---|---|---|
| | | **N2** | | **N1** | | **p2** | | **p1** | | **k1** | | **k2** | | **N0** | |
| | **cell** | **dark** | **light** | **dark** | **light** | **dark** | **light** | **dark** | **light** | **dark** | **light** | **dark** | **light** | **dark** | **light** |
| Fit to individual traces | 1 | 63.2549 | 4.0891 | 1014.821 | 612.9435 | 0.6552 | 1 | 0.0381 | 0.036 | 6.7047 | 3.8195 | 0.1131 | 0.0565 | 10951.11 | 6108.06 |
| | 2 | 88.8845 | 24.3924 | 1119.725 | 481.5995 | 0.4589 | 0.7038 | 0.0198 | 0.0262 | 3.6896 | 6.8613 | 0.0979 | 0.1179 | 15075.5 | 5136.241 |
| | 3 | 43.6184 | 14.9597 | 1193.699 | 208.3005 | 0.4844 | 0.483 | 0.0245 | 0.0401 | 3.0565 | 7.0364 | 0.2364 | 0.1792 | 4858.391 | 2705.826 |
| | 4 | 29.1179 | | 871.7712 | | 0.6498 | | 0.0232 | | 4.799 | | 0.0616 | | 12952.08 | |
| | 5 | 51.9496 | | 680.1761 | | 0.7939 | | 0.044 | | 5.7975 | | 0.4998 | | 4233.132 | |
| | 6 | 62.8848 | 40.9617 | 756.2691 | 462.0405 | 0.5731 | 0.5121 | 0.0399 | 0.0388 | 6.6169 | 4.4341 | 0.1757 | 0.0636 | 5934.726 | 4183.475 |
| | 7 | 93.5481 | | 612.3086 | | 0.602 | | 0.0512 | | 7.8715 | | 0.3266 | | 7862.92 | |
| | 8 | 67.5227 | 37.1755 | 792.4491 | 184.7107 | 0.5939 | 0.573 | 0.0371 | 0.0539 | 3.4662 | 6.3637 | 0.1172 | 0.3467 | 16124.21 | 3107.672 |
| | 9 | 140.5941 | | 479.3769 | | 0.3925 | | 0.0367 | | 6.8927 | | 0.2926 | | 6043.896 | |
| | 10 | 65.6747 | 90.2895 | 630.9655 | 579.8763 | 0.6345 | 0.6507 | 0.0468 | 0.074 | 4.9007 | 4.5641 | 0.1065 | 0.0297 | 33308.25 | 3568.922 |
| | mean | 70.705 | 35.3113 | 815.1562 | 421.5785 | 0.5838 | 0.6538 | 0.0361 | 0.0448 | 5.3795 | 5.5132 | 0.2027 | 0.1323 | 11734.4215 | 4135.0327 |
| | SEM | 9.8341 | 12.3373 | 73.7226 | 74.9472 | 0.0363 | 0.0771 | 3.32E-03 | 6.87E-03 | 0.5211 | 0.5714 | 0.0433 | 0.0481 | 2753.8168 | 526.5037 |
| | median | 64.4648 | 30.784 | 774.3591 | 471.82 | 0.5979 | 0.6118 | 0.0376 | 0.0394 | 5.3491 | 5.4639 | 0.1465 | 0.0908 | 9407.015 | 3876.1985 |
| | Mann-Whitney U-test (p value) | 0.034 | | 0.006 | | 0.625 | | 0.357 | | 0.871 | | 0.357 | | 0.011 | |
| | Bonferroni adjusted (p value) | 0.238 | | 0.042 | | >0.99 | | >0.99 | | >0.99 | | >0.99 | | 0.077 | |
| Fit to grand avarage | | 68.1721 | 18.8849 | 798.3338 | 229.7779 | 0.5747 | 0.6633 | 0.035 | 0.0513 | 5.1283 | 5.4453 | 0.167 | 0.0544 | 7948.425 | 2248.991 |
